# SARS-CoV-2 Spike Protein Accumulation in the Skull-Meninges-Brain Axis: Potential Implications for Long-Term Neurological Complications in post-COVID-19

**DOI:** 10.1101/2023.04.04.535604

**Authors:** Zhouyi Rong, Hongcheng Mai, Saketh Kapoor, Victor G. Puelles, Jan Czogalla, Julia Schädler, Jessica Vering, Claire Delbridge, Hanno Steinke, Hannah Frenzel, Katja Schmidt, Özüm Sehnaz Caliskan, Jochen Martin Wettengel, Fatma Cherif, Mayar Ali, Zeynep Ilgin Kolabas, Selin Ulukaya, Izabela Horvath, Shan Zhao, Natalie Krahmer, Sabina Tahirovic, Ali Önder Yildirim, Tobias B. Huber, Benjamin Ondruschka, Ingo Bechmann, Gregor Ebert, Ulrike Protzer, Harsharan Singh Bhatia, Farida Hellal, Ali Ertürk

## Abstract

Coronavirus disease 2019 (COVID-19), caused by the severe acute respiratory syndrome coronavirus type 2 (SARS-CoV-2), has been associated mainly with a range of neurological symptoms, including brain fog and brain tissue loss, raising concerns about the virus’s acute and potential chronic impact on the central nervous system. In this study, we utilized mouse models and human post-mortem tissues to investigate the presence and distribution of the SARS-CoV-2 spike protein in the skull-meninges-brain axis. Our results revealed the accumulation of the spike protein in the skull marrow, brain meninges, and brain parenchyma. The injection of the spike protein alone caused cell death in the brain, highlighting a direct effect on brain tissue. Furthermore, we observed the presence of spike protein in the skull of deceased long after their COVID-19 infection, suggesting that the spike’s persistence may contribute to long-term neurological symptoms. The spike protein was associated with neutrophil-related pathways and dysregulation of the proteins involved in the PI3K-AKT as well as complement and coagulation pathway. Overall, our findings suggest that SARS-CoV-2 spike protein trafficking from CNS borders into the brain parenchyma and identified differentially regulated pathways may present insights into mechanisms underlying immediate and long-term consequences of SARS-CoV-2 and present diagnostic and therapeutic opportunities.

**Graphical Summary:** 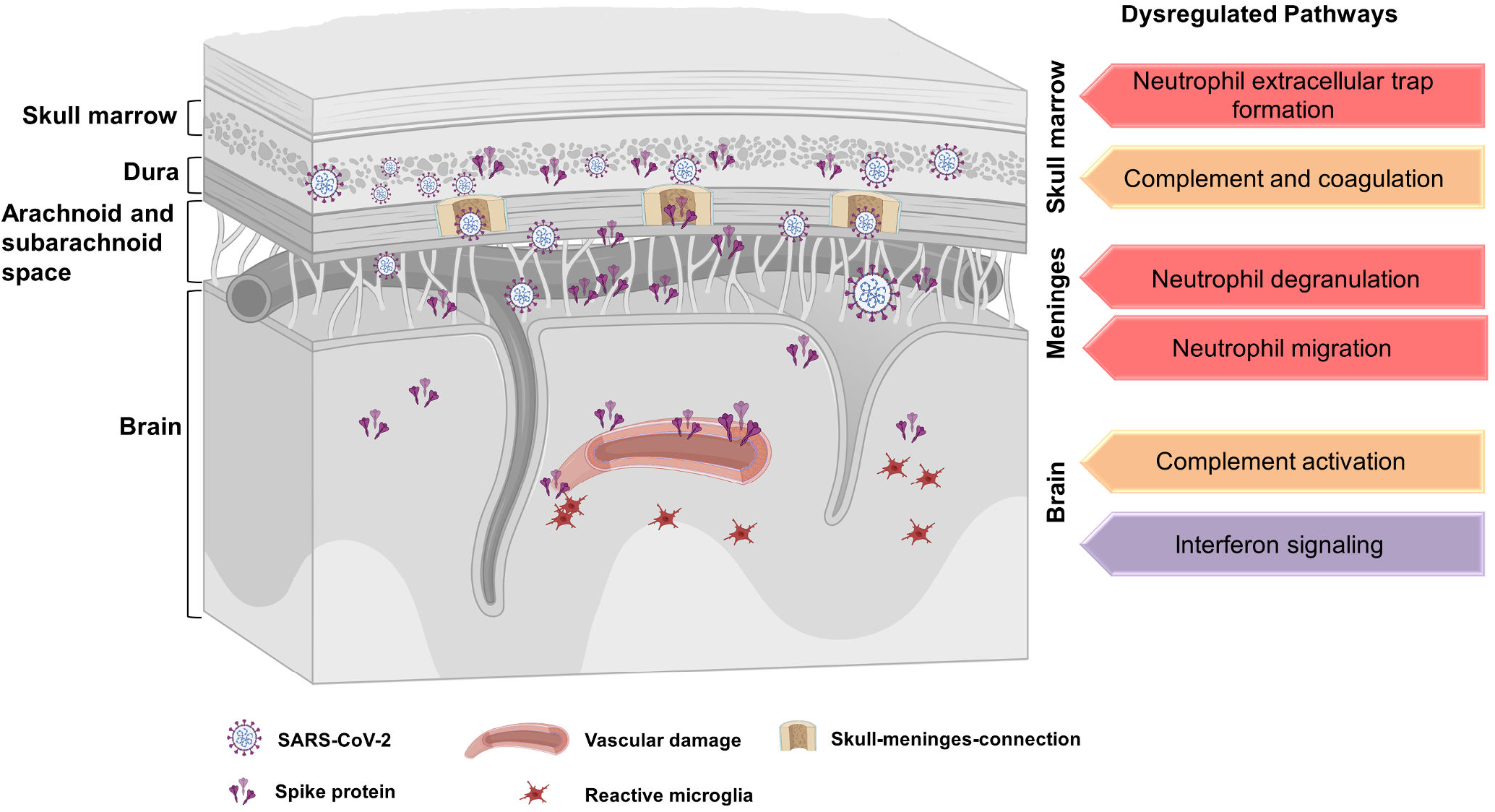

**Short Summary:** **The accumulation of SARS-CoV-2 spike protein in the skull-meninges-brain axis presents potential molecular mechanisms and therapeutic targets for neurological complications in long-COVID-19 patients**.

## INTRODUCTION

SARS-CoV-2 infection is associated with numerous neurological and neuropsychiatric complications^1-3^, including anosmia, dysgeusia, fatigue, myalgia, depression, headache, encephalopathy and meningitis and also substantially increase the risk for ischemic strokes^4,5^. Even patients with mild cases of COVID-19 often suffer from long-term SARS-CoV-2 effects in the brain, including fogging, reduced grey matter thickness, and brain size^6–8^. Several studies have investigated the involvement of the central nervous system (CNS) in COVID-19-related symptoms, and although SARS-CoV-2 was detected in brain tissue in some samples and studies^3,9–12^, other studies failed to detect the virus^12–16^. Various technical issues such as contamination from the blood in PCR-based methods, misidentification of capillaries as parenchyma in immunohistochemistry, staining errors using inappropriate antibodies^17^ or differences in patient populations might explain this discrepancy. However, even without detectable virus RNA in the brain parenchyma, signs of widespread immune activation could be detected^18^. The lack of evidence for the viral presence and especially viral replication in the brain led to the hypothesis that virus-shed proteins circulating in the bloodstream may promote an inflammatory response independent of direct viral infection of the affected organs, including the brain^19,20^. Notably, the highly immunogenic spike protein, also used in COVID-19 vaccines^21–23^, might be a candidate for triggering infection-independent effects.

The spike protein has been shown to affect endothelial function *in vitro*^24–26^ and *in vivo*^27,28^ and induce TLR2-mediated inflammatory responses in vitro after intraperitoneal injection in mice^29^, but whether such responses can also be observed in patients has not been thoroughly investigated. However, the long persistence of the spike protein has been shown in the patient’s immune cells (at least 15 months)^30^ and in the patient’s blood plasma (at least 12 months in a preprint)^31^. Radio-labeled free spike protein has been shown to cross mice’s blood-brain barrier and enter the brain parenchyma^32^. However, due to the limited resolution of the methods employed, the exact routes of spike protein entry to the brain, their targets, and molecular changes associated with spike protein accumulation in brain tissue remain largely unclear^33,34^.

Here, we used optical tissue clearing to identify all tissues that accumulated SARS-CoV-2 spike protein in mice and investigated the distribution of spike protein in post-mortem samples from COVID-19 patients. We also characterize the protein expression consequences of SARS-CoV-2 infections in different skull tissues from post-mortem human samples with mass spectrometry-based proteomics. We found an accumulation of spike protein in the skull marrow niches, recently discovered skull-meninges connection (SMC)^35–40^, meninges, and the brain parenchyma in both mouse and human samples. The human proteomics data showed dysregulation of complement and coagulation cascades, neutrophil-related pathways, and an upregulation of pro-inflammatory proteins. Injecting spike protein to skull marrow niches directly in healthy mice triggered proteome changes and cell death in the brain parenchyma. Surprisingly, we identified lingering spike protein in the skull samples of a subset of individuals who recovered from COVID-19 and died due to non-COVID-related causes. Thus, accumulation of SARS-CoV-2 and spike protein at the CNS borders can contribute to changes in the brain, suggesting a possible mechanism for the neurological effects of SARS-CoV-2 infection.

## RESULTS

### Whole body distribution of spike S1 protein in a mouse model

To identify all potential tissues that are targeted by SARS-CoV-2, we mapped the distribution of fluorescently labeled spike S1 protein in intact transparent mice. While not a direct measurement of viral tropism, recent studies demonstrated the utility of the recombinant spike protein as a proxy to study SARS-CoV-2 targeting and pathology in mice^32,27^.

WT spike protein has a low binding affinity for the mouse angiotensin-converting enzyme 2 (ACE2)^41,42^. We could not use the transgenic K18-hACE2 mouse line, which arbors the human form of ACE2 (hACE2) driven under the cytokeratin 18 promoter leading to an ectopic overexpression of hACE2^43^ which would bias the biodistribution analysis. We, therefore, conjugated Alexa-647 to spike S1 (N501Y) protein carrying the mutation, which enables binding to the mouse ACE2 isoform. As a control, we also conjugated Alexa-647 to the WT spike and influenza virus hemagglutinin (HA) for comparison. At 30 minutes post-intravenous injection, the mice were transcardially perfused, and their whole bodies were subjected to optical transparency clearing and imaged using light-sheet microscopy^44^. We detected spike S1 binding in most organs, including the heart, lung, liver, kidney, intestine, thymus, spleen, and pancreas, in contrast to the WT spike S1 and HA more exclusive to the liver and lung (**Fig. 1A**). Within each organ, the spike S1 highly accumulated in close vicinity to blood vessels (**Fig 1B**, arrowheads vs. arrows) in the liver, kidney, and lung (**Video 1**). Most of the spike S1 signal in the abdomen was situated in the capillary bed and most likely from liver Kupffer cells, spleen follicles, glomeruli, and alveoli. Spike S1 protein was also found in the parenchyma of the testis and ovary (**Fig. S1A, Video 2**). The multi-organ distribution of spike S1 binding is consistent with the known expression pattern of ACE2, widely expressed in various organs and serving as the cell entry receptor for SARS-CoV-2^45,46^.

**Figure 1.**
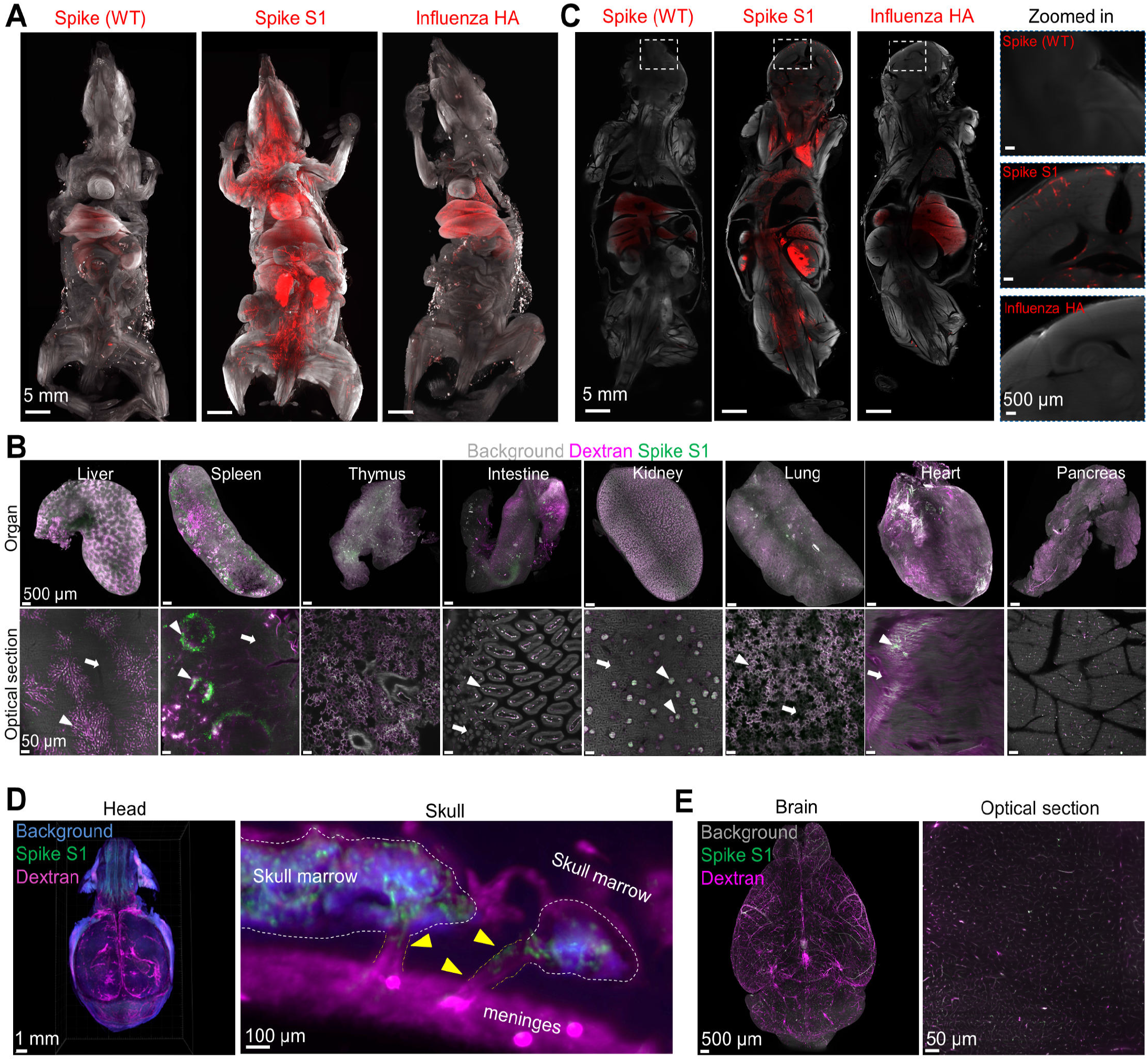
Spike protein exhibits multi-organ binding capacity. (A) 3D reconstructions of whole mouse body after spike protein (WT), spike S1 (N501Y), and HA injection, imaged with light-sheet microscopy. (B) 3D reconstruction of main internal organs and representative high-resolution optical section view with spike S1 (N501Y) protein and dextran labeled vasculature. Arrow heads and arrows indicate regions with and without spike S1 protein, respectively. (C) Optical section of whole-body images after spike protein (WT), spike S1 (N501Y), and HA injection. The white box area was zoomed in to check the brain. (D) Visualization of spike S1 (N501Y) protein in the intact mouse head and representative sagittal images of the skull bone marrow, SMCs and meninges. Arrow heads indicate spike S1 protein in SMCs. (E) Representative images of spike S1 (N501Y) protein in the brain.

Moreover, spike S1 was also detected in the brain prefrontal cortex (**Fig. 1C, Video 3**). Further examining the mouse heads, we found substantial spike S1 accumulation in the skull marrow niches (**Fig. 1D, Video 4**). Notably, we detected the spike protein in the channels connecting the skull marrow to the meninges (**Fig. 1D**), suggesting that the SMCs could play a role in distributing the S1 protein through the skull in addition to other potential routes, such as transport by phagocytic cells or direct extravasation from blood vessels. Similarly, the spike S1 protein accumulated in the marrow from other bones like the long bones, tibia, and femur, indicating it reaches bone marrow niches (**Fig. S1B**). In the CNS, we found spike protein in the diverse regions of the brain and spinal cord (**Video 5**). Co-staining with the dextran for vessel visualization, we found that the spike S1 protein was mainly localized in the brain blood vessels (**Fig. 1E**)

### SARS-CoV-2 infection in the human skull, meninges, and brain

The skull bone marrow was characterized as myeloid cell reservoirs for the meninges and CNS parenchyma through the channels between skull marrow and meninges^35–37,47^. The lymphopoietic niche is also identified in the CNS border meninges^38–40^, suggesting the involvement of the skull and meninges in neurological diseases. One study reported the transition of bacteria from the cerebrospinal fluid through the channels into skull marrow cavities^48^. Therefore, we hypothesized that the connections between skull marrow and meninges^49–54^ might contribute to SARS-CoV-2 virus entry into the brain. We first studied the distribution of the SARS-CoV-2 in post-mortem skull tissues of patients who died from COVID-19 infection (n = 17 different COVID-19 cases, n = 10 had also meninges attached, **Fig. 2A, Table 1**). We used the optical clearing SHANEL protocol developed for large human tissue^55^ and three-dimensional (3D) imaging to analyze the histopathological profile of human skull samples with the attached dura mater. We identified viral spike protein in the skull marrow niches, SMCs, and meninges of SARS-CoV-2 infected patients (**Fig. 2B, Video 6 and 7**). There was no spike protein labeling in samples from control individuals (**Fig. S2A**). We found spike protein in collagen IV and CD31-positive blood vessels but also 44.8% outside of the blood vessels in the skull marrow niches (**Fig. 2C and 2D**), suggesting that spike protein leaked into skull marrow (Video 8 and 9). Furthermore, with confocal imaging, we localized spike protein in the perinuclear space of meningeal cells and the vicinity of NeuN-positive neurons in the brain cortex (**Fig. 2E**). We also analyzed the lungs of these patients, where we identified co-expression of ACE2 and spike protein (**Fig. S2B**). In the meninges of infected patients, we found a concomitant decrease of ACE2 (**Fig. S2C**) consistent with reported ACE2 downregulation and endothelial dysfunction in COVID-19 patients^28^.

**Figure 2.**
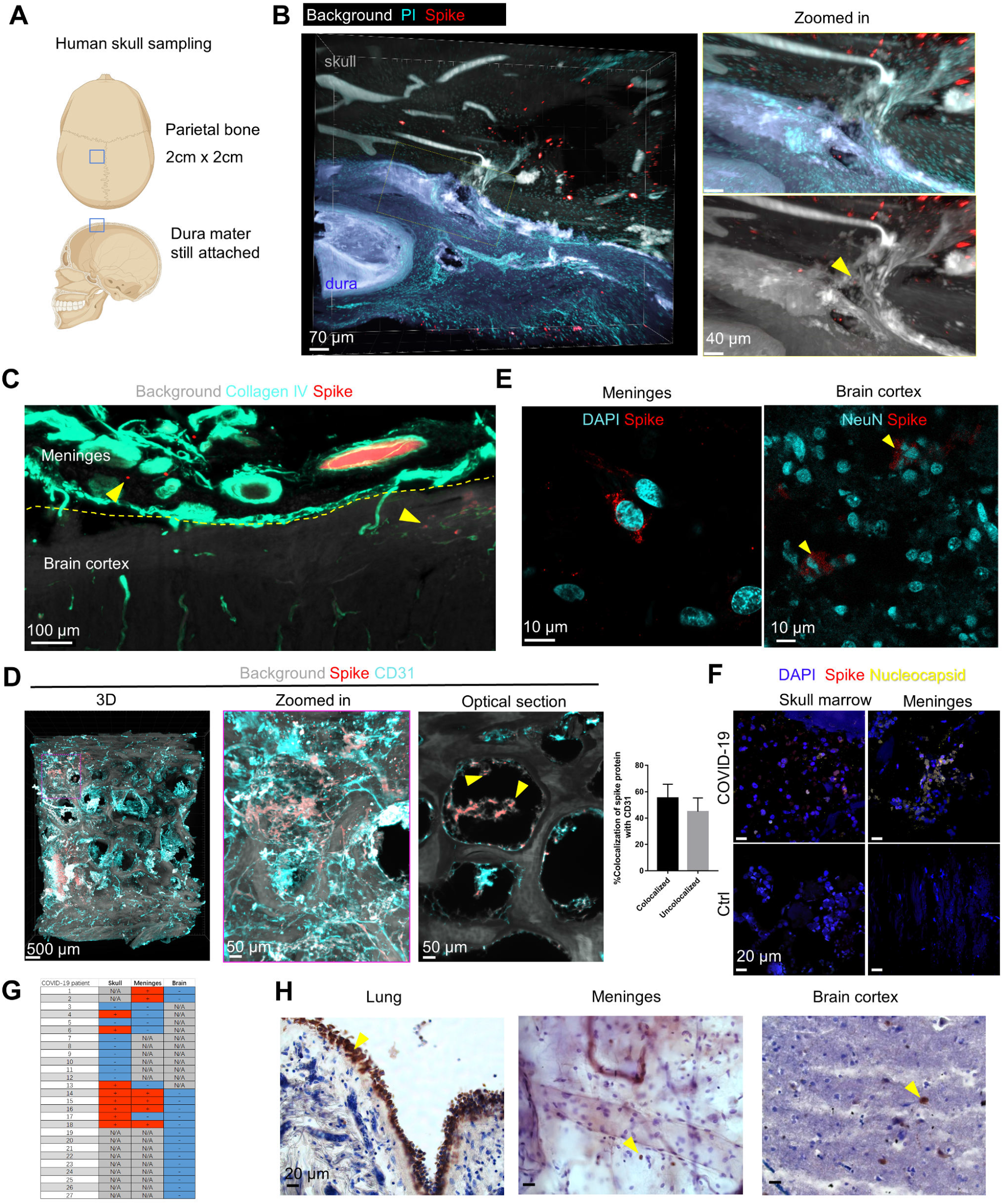
SARS-CoV-2 infection in the human skull, meninges, and brain. (A) Illustration for the sampling of human skull with meninges. (B) Representative images of spike protein antibody and PI labeling in the COVID-19 patient skull with meninges. The dura mater (blue) is segmented manually based on autofluorescence and imported for 3D reconstruction in Imaris. The yellow arrowhead indicates positive signal in the SMC. (C) Representative image of spike protein and collagen IV antibody staining in COVID-19 patient brain with meninges. (D) Representative images of spike protein antibody and CD31 antibody labeling in the COVID-19 patient skull with meninges. Arrowheads indicate spike protein in skull marrow niche. Quantification of spike protein colocalization with CD31 signal in 6 optical sections. Data are mean ± SEM. (E) Representative confocal images of spike protein antibody labeling in the COVID-19 patient meninges and brain cortex. (F) Representative confocal images of spike protein and nucleocapsid protein in human skull marrow and meninges. (G) RT-PCR results of COVID-19 patient skull, meninges, and brain samples. N/A, not available. (H) Immunohistochemistry of spike protein in COVID-19 patient lung, meninges, and brain cortex. Spike protein shown in brown, cell nucleus in blue. Yellow arrows indicate positive regions.

SARS-CoV-2 RNA was detected in 8/16 skull samples and 6/12 meninges samples of COVID-19 patients using RT-PCR, along with the nucleocapsid protein (**Fig. 2F**). All the brain prefrontal cortex samples we inspected were PCR negative (**Fig. 2G**), and we also did not detect the nucleocapsid protein (**Fig. S2D**). The presence of spike protein in the absence of viral load (PCR negative) in the brain and meninges from the same patients (**Fig. 2H**) would suggest either a specific uptake mechanism to the brain or a longer half-life of spike protein compared to SARS-CoV-2 viral particles.

### Proteomics profiling of COVID-19 patient skull marrow, meninges, and brain samples

To explore the consequences of SARS-CoV-2 infection and spike protein accumulation at the CNS borders, we performed mass spectrometry-based label-free quantitative proteomics analysis of region-matched human tissues from COVID-19 patients and controls (non-COVID-19) (**Table 1**).

In the skull marrow samples, a total of 6504 proteins m were identified across the samples from COVID-19 patients (n = 10) and controls (n = 10) (only samples with both skull and meninges from the same patients were analyzed, **Fig. S3A**). While we observed high concordance within the groups (Pearson correlation coefficients: 0.67 to 0.87 in controls and 0.68 to 0.93 in COVID-19 samples) (**Fig. S3B**), the correlation between COVID-19 and control samples was relatively low to moderate (Pearson correlation coefficients: 0.55 to 0.84) suggesting a substantial and reproducible influence of the viral infection on the protein expression profiles in the skull marrow. The principal component analysis (PCA) plot showed clear segregation of the skull marrow samples from COVID-19 and control group (**Fig. 3A**). We identified 655 differentially expressed proteins in the COVID-19 skull marrow, with 496 being upregulated and 159 being downregulated (**Fig. S3C**). We found a reduced expression of SARS-CoV-2 host cell-associated entry factors neuropilin 1 (NRP1)^56^. However, other coronavirus entry factors such as neuropilin 2 (NRP2), dipeptidyl peptidase 4 (DPP4, also known as CD26)^57,58^, basignin (BSG, also known as CD147), and alanyl aminopeptidase (ANPEP, also known as CD13)^57^, and cathepsin B (CTSB)^59^ remained unchanged.

**Figure 3.**
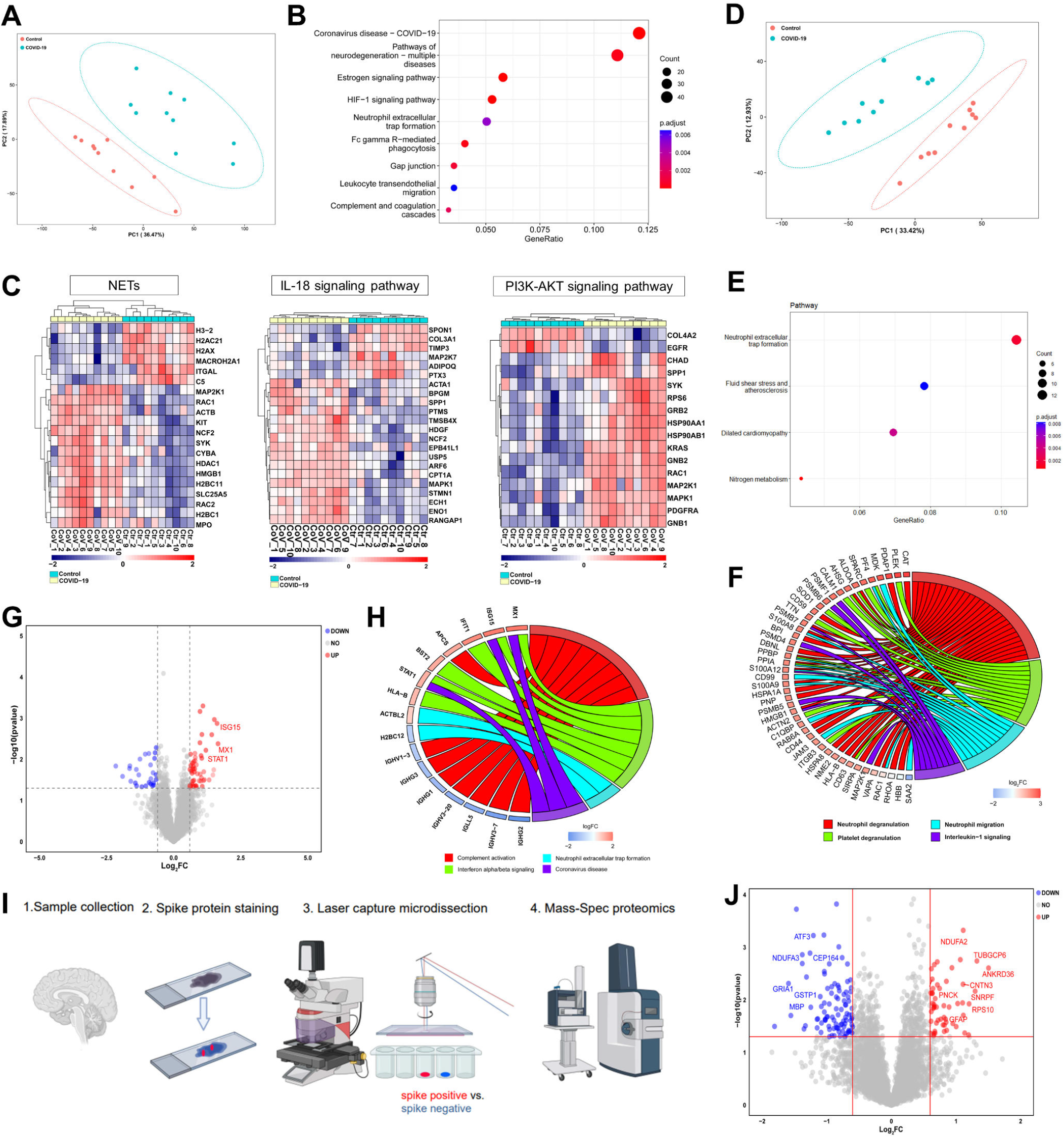
Proteomics profiling of COVID-19 patient skull, meninges, and brain samples. (A) PCA plot between COVID-19 and non-COVID-19 control skull marrow. n = 10 biological replicates. (B) KEGG pathway analysis showing the most dysregulated pathway in COVID-19 skull marrow compared to non-COVID-19 control. (C) The dysregulated proteins related to neutrophil extracellular traps (NETs), IL-18 signaling pathway and PI3K-AKT signaling pathway in skull marrow are listed in heatmaps. (D) PCA plot between COVID-19 and control meninges (p < 0.05, Log2FC = 1.5). n = 10 biological replicates. (E) KEGG pathway analysis showing the most dysregulated pathway in COVID-19 meninges compared to non-COVID-19 control. (F) Chord diagram showing the most enriched biological processes with their differentially expressed proteins in the COV-ID-19 meninges. (G) Volcano plot showing differentially expressed genes between COVID-19 and control brain cortex samples (p < 0.05, Log2FC = 1). n = 11 COVID-19 and 8 non-COVID-19 control samples. FC, fold change. (H) Chord diagram showing the most enriched biological processes with their differentially expressed proteins in the COVID-19 brain cortex. (I) Illustration of proteomics analysis of spike protein-positive and -negative brain regions from COVID-19 patients after laser capture microdissection. (J) Volcano plot showing differentially expressed genes between spike protein positive and negative regions (p < 0.05, Log2FC = 0.6). n = 3 COVID-19 patients. FC, fold change.

We used gene set enrichment analysis to correlate the dysregulated proteins to biological processes modulated in response to SARS-CoV-2 infection. The majority of the proteins annotated as involved in the coronavirus disease pathway in the KEGG database were also upregulated in the skull marrow of COVID-19 patients in our data, which confirms the effects of SARS-CoV-2 infection in the skull marrow of COVID-19 patients (**Fig. 3B**). We also detected upregulation of interleukin enhancer-binding factor 3 (ILF3) which has been previously associated with viral replication^60^. The most strongly dysregulated part of the immune system was the complement system, which was previously reported to be activated in COVID-19^61^. High-severity infection of COVID-19 is often associated with hyperactivation followed by exhaustion of complement proteins such as C3 and C4^62,63^. Indeed we found significant downregulation of complement components C1, C4, C5, C8, and factor H in the skull marrow of these non-survivor COVID-19 patients and decreased coagulation factor XI expression. Beyond the complement system, we also identified upregulation of critical regulators of interferon-alpha/beta signaling cascades such as interferon-induced GTP-binding protein Mx1 (MX1), interferon-stimulated gene 20 kDa protein (ISG20) as well as signal transducer and activator of transcription 1-alpha/beta (STAT1), an essential regulator of IL-6 signaling in the skull marrow (**Fig. S3D**). Such an inflammatory response is responsible for the cytokine storms described in COVID-19, further associated with increased accumulation and degranulation of neutrophils^64,65^. De facto, some of the proteins involved in neutrophil degranulation, such as high mobility group protein B1 (HMGB1), and heat shock protein HSP 90-alpha (HSP90AA1), were also upregulated in the skull marrow of COVID-19. Furthermore, we found expression changes in proteins related to the IL-18^66^ and PI3K-AKT signaling pathways^67^, which have been implicated in SARS-CoV-2 infections and are associated with coagulopathies, for example, upregulation of HSP90AA1, HSP90AB1 and GNB1 and downregulation of EGFR (**Fig. 3C**).

In the human meninges’ proteomics, we identified 4927 proteins across all samples (**Fig. S3E**). The PCA plot clearly distinguishes the meninges samples from COVID-19 and control groups (**Fig. 3D**). Here, we identified 373 differentially expressed proteins with 349 upregulated and 24 downregulated (**Fig. S3F**). Our pathway analysis revealed that proteins involved in the neutrophil degranulation were significantly upregulated in the meninges of COVID-19 as compared to controls (**Fig. 3E**). These include PDAP1, DBNL, CD44, RAB6A, and HMGB1 (**Fig. 3F**). These proteins could be linked to the formation of NETs in response to the virus. As in the COVID-19 skull marrow niches, we observed upregulation of HMGB1, an extracellular protein that has been shown to stimulate NETs formation^68^. PI3K-AKT pathway proteins were also dysregulated in the meninges (**Fig. 3F**). HSP90AA1 was upregulated in the COVID-19 skull and meninges. We also observed the overexpression of calprotectin (S100A8/A9) in the meninges, which plays an essential pro-inflammatory role in the migration of neutrophils (**Fig. 3F**). Specifically, S100A8 has recently been hypothesized to be involved in hyper-inflammation in severe COVID-19^69^. We also identified upregulation of platelet factor 4 (PF4) and Platelet basic protein (PPBP) in the meninges tissue (**Fig. S3G**).

These data suggest that neutrophils, which constitute the major cell population in the skull marrow niches^70^, might move into the meninges, induce the NETs formation, and trigger pro-inflammatory responses. Indeed, neutrophil migration directly from vascular channels connecting skull marrow to the meninges has been reported in inflammatory events^35,36^.

We dissected samples from the brain cortex of 8 COVID-19 patients and 11 control cases for proteomics analysis and identified 7138 proteins. Of these, 76 proteins were differentially expressed, with 49 upregulated and 27 downregulated (**Fig. 3G**). The predominant dysregulated pathways were complement activation, NETs formation, coronavirus disease, and the interferon-alpha/beta signaling pathways (**Fig. 3H**). We identified PYCARD and NLRC3 as upregulated proteins previously associated with assembling inflammasome^71,72^. Among the other upregulated proteins, we also identified BST2, which restricts the viruses from infecting adjacent cells. In addition, we also investigated COVID-19 brain tissue alterations and analyzed microgliosis in COVID-19 brain tissue compared to controls. We identified activated myeloid cells with enlarged cell body morphologies (**Fig. S3H**)^73^. This data is in line with our proteomics data suggesting a persistent inflammatory state in the brain of COVID-19 patients. Finally, we found micro-bleedings in all four COVID-19 brain tissues that we analyzed using Prussian blue staining, while in only one of the four brain tissues in the control group (**Fig. S3I**).

A comparison of datasets from the skull marrow, meninges, and brain samples identified five common differentially expressed proteins related to antigen presentation and recognition, including IGHV1-3, HLA-B, APCS, ISG15, and MX1, whereas MX1 is known to have an antiviral function^74^.

To further pinpoint the consequences of spike protein-specific effects in the brain tissue beyond the acute inflammatory response, we performed targeted proteomics on laser capture micro-dissected tissues^75^ by analyzing the spike positive vs. spike negative brain regions (**Fig. 3I**). We identified several dysregulated proteins associated with neurodegeneration, such as downregulation of MBP and upregulation of GFAP. GFAP has been described as a biomarker to detect damage in the blood-brain barrier leading to brain injury during COVID-19 infection^76^. GRIA1 is a glutamate receptor subunit associated with neurodevelopmental disorders, and its deletion in mice causes attention deficit and sleep disorder, both reported symptoms of post-COVID syndrome^77–80^. NDUFA3 is a subunit of the mitochondrial membrane respiratory chain associated with Alzheimer’s disease^81^. We found NDUFA2 and NDUFA3 were dysregulated in the spike protein-positive regions (**Fig. 3J**). To our knowledge, their role in COVID-19 pathogenesis has not been investigated in detail yet. However, their decrease or loss of function significantly impairs mitochondrial function^82^, a source of oxidative stress also reported in SARS-CoV-2 infection^83^ and long-term symptoms such as chronic fatigue^84^.

### Spike S1 protein triggers proteomics changes in the mouse skull marrow, meninges, and brain

To investigate the potential impact of spike S1 protein binding in the skull marrow, meninges, and brain tissues without other viral proteins, we performed proteomics analysis of these three areas at three days post intravenous injection of spike S1 (N501Y), HA protein, and saline controls in WT mice.

We identified 9167 proteins across the skull marrow samples from spike S1, HA, and control groups (**Fig. S4A**). The PCA plot showed skull marrow samples after spike S1 injection separated distinctly from the control group. In contrast, the HA group partially overlapped with the control (**Fig. 4A**). Compared to the saline control, we identified 808 differentially expressed proteins in the skull marrow after spike S1 injection, with 348 upregulated and 460 being downregulated (**Fig. S4B**). While in the skull marrow after HA injection, we only identified 16 differentially expressed proteins compared to the control (**Fig. S4C**). We found a substantial differential expression in ribosomal proteins and proteins involved in NETs formation, neutrophil degranulation, and PI3K-AKT pathways (**Fig. 4B and 4C**), which were also significantly dysregulated in COVID-19 patient skulls.

**Figure 4.**
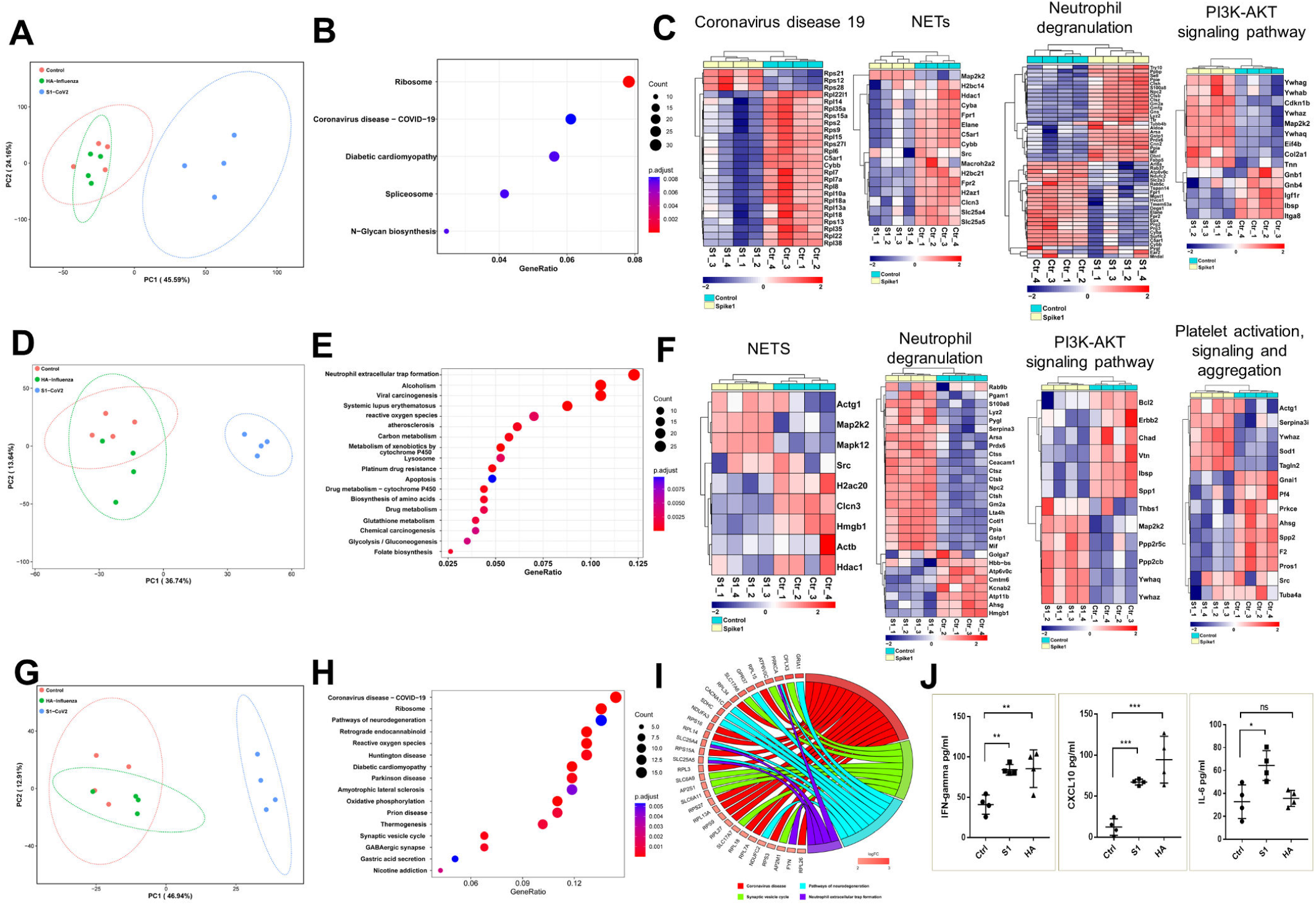
Spike protein triggers proteomics changes in the mouse skull, meninges, and brain. (A) PCA plot of skull marrow samples. (B) KEGG pathway analysis showing the most dysregulated pathways in skull marrow after spike S1 injection compared to control. (C) The dysregulated proteins related to various pathways in skull marrow after spike S1 injection are listed in heatmaps. (D) PCA plot of meninges samples. (E) KEGG pathway analysis showing the most dysregulated pathways in meninges after spike S1 injection compared to control. (F) The dysregulated proteins related to various pathways in meninges after spike S1 injection are listed in heatmaps. (G) PCA plot of brain samples. (H) KEGG pathway analysis showing the most dysregulated pathways in skull marrow after spike S1 injection compared to control. (I) Chord diagram showing the most enriched biological processes with their differentially expressed proteins in the brain after spike S1 injection. (J) Plasma ELISA of IFN-gamma, CXCL10, and IL-6 at 3 days post injection. Graph displays mean ± SEM. Two-tailed unpaired Student’s t test, * p < 0.05; ** p < 0.01; *** p < 0.001; ns, nonsignificant.

Similarly, we identified 8687 proteins across the meninges samples from spike S1, HA, and control groups (**Fig. S4D**). After spike S1 injection, the meninges samples distinctly separated from the control group in the PCA plot, whereas the HA group partially overlapped with the control (**Fig. 4D**). We identified 426 differentially expressed proteins in the meninges after spike S1 injection, with 241 being upregulated and 185 being downregulated (**Fig. S4E**). Pathway analysis suggested the most abundant dysregulation in the NETs formation. We also found proteins enriched in biological processes, including neutrophil degranulation, PI3K-AKT signaling pathway, and platelet activation, signaling, and aggregation (**Fig. 4E and 4F**). While in the meninges after HA injection, we only identified 33 differentially expressed proteins compared to the control (**Fig. S4F**). After spike S1, HA, and saline intravenous injection, we identified 8104 proteins in the brain cortex (**Fig. S4G**). The brain cortex samples after spike S1 injection also showed a clear segregation from the HA and control group in the PCA plot (**Fig. 4G**). 213 proteins were identified to be dysregulated in the brain cortex after spike S1 injection, with 190 being upregulated and 23 being downregulated (**Fig. S4H**). In addition to the altered level of ribosomal proteins associated with COVID-19 in the spike S1 group compared to the control group, we identified proteins dysregulated in pathways of neurodegeneration (**Fig. 4H and 4I**). GRIA1 and NDUFA3 were also dysregulated in the spike protein-positive COVID-19 patient brain regions (**Fig. 3J**). Whereas in the brain cortex after HA injection, only one protein STT3A, a catalytic subunit of the oligosaccharyl transferase complex, was identified to be dysregulated (**Fig. S4I**).

On the other hand, we tested the plasma cytokine levels of the mice three days after injection of spike S1, HA, and saline control. The spike S1 and HA protein caused increased inflammation, as shown by IFN-gamma and CXCL10 enzyme-linked immunosorbent assay (ELISA). We also noticed a difference in spike S1 and HA protein-induced cytokine levels, and plasma IL-6 was increased three days after spike S1 injection but not changed after HA injection (**Fig. 4J**).

These data suggest that the SARS-CoV-2 spike S1 protein alone is sufficient to trigger a broad proteomic change in the skull marrow, meninges, and brain compared to the HA protein. The dysregulated proteins are enriched in pathways related to coronavirus disease, NETs formation, and neutrophil degranulation, consistent with the changes observed in virus-infected human samples. Our data also suggest that spike S1 protein circulation in the blood would be an important trigger of COVID-19-related broader proteome change in the skull marrow, meninges, and brain cortex.

### Spike protein from the skull marrow leads to brain cortex neuronal injury

To understand whether the spike protein could be released from the skull marrow niches and induce acute and long-term pathological changes in the brain parenchyma, we injected the spike S1 (N501Y) directly into the skull marrow (**Fig. 5A and 5B**)^35^. We found that the spike S1 protein reached the meninges and brain parenchyma 30 minutes after microinjection to the skull (**Fig. 5C**).

**Figure 5.**
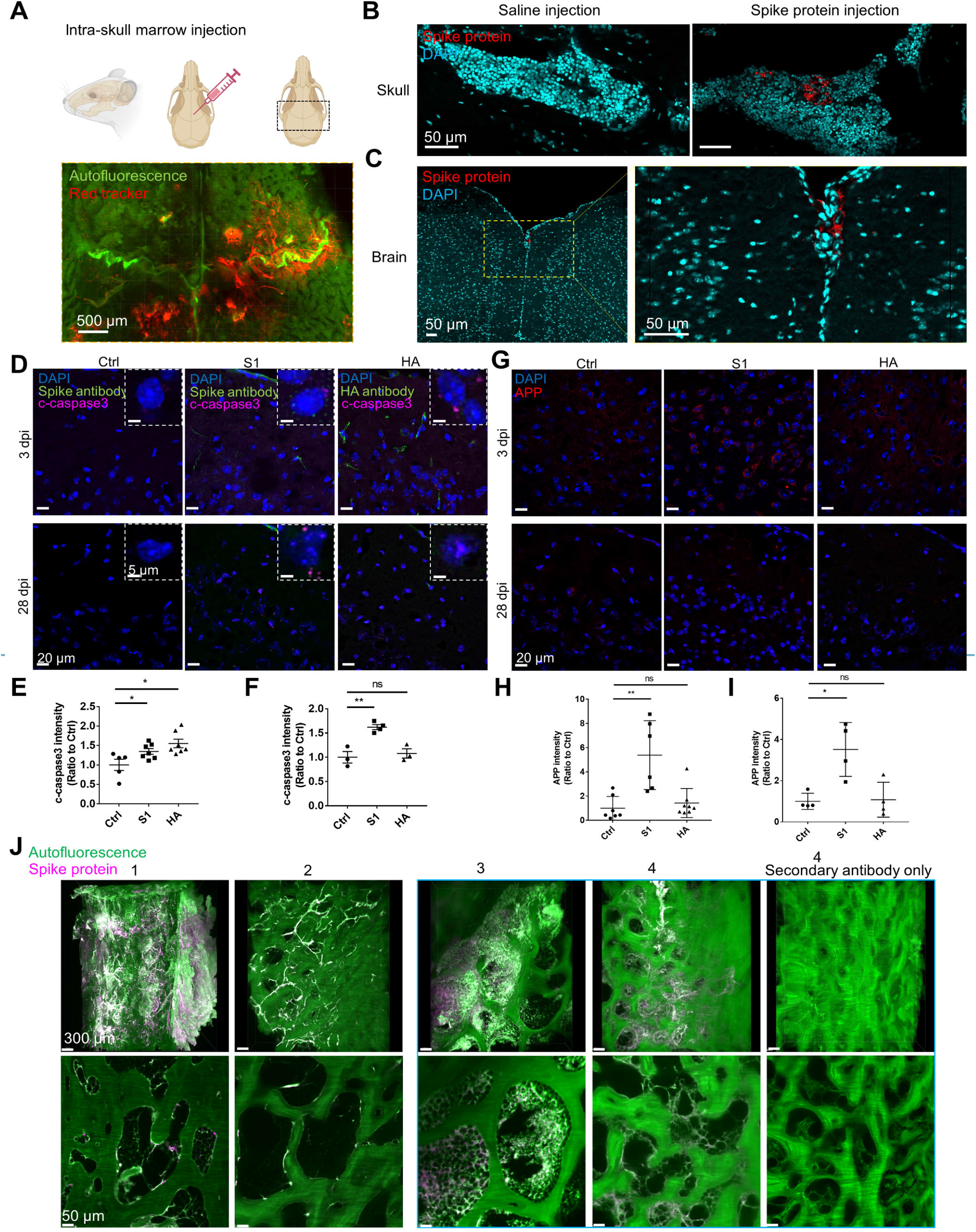
Spike protein from the skull marrow leads to brain cortex neuronal injury. (A) Illustration for intra-skull microinjection. (B) Representative images of spike protein antibody and DAPI labeling in the mouse skull 30 minutes after intra-skull injection of spike S1 protein and saline control. (C) Representative images of spike protein antibody and DAPI labeling in the mouse brain cortex 30 minutes after intra-skull injection of spike S1 protein. (D) Representative images of c-caspase3 antibody, spike antibody, and HA antibody staining in mouse brain cortex after intra-skull injection of spike S1, HA protein, and saline control. (E) Quantification of the c-caspase3 intensity in brain cortex 3 days post injection, n = 3. (F) Quantification of the c-caspase3 intensity in brain cortex 28 days post injection, n = 3. (G) Representative images of APP antibody staining in mouse brain cortex after intra-skull injection of spike S1, HA protein, and saline control. (H) Quantification of the APP intensity in brain cortex 3 days post injection, n = 3. (I) Quantification of the APP intensity in brain cortex 28 days post injection, n = 3. (J) Representative SHANEL tissue clearing and imaging of human skull samples with spike protein antibody staining. A Z-stack of 1200 μm thick consecutive sections were reconstructed in 3D with Imaris. Patient 1, confirmed fatal case of COVID-19; Patient 2, died before COVID-19 pandemic; Patient 3 and 4, died during COVID-19 pandemic, cause of death is not COVID-19. Data are mean ± SEM. Two-tailed unpaired Student’s t test. * p < 0.05. ** p < 0.01; ns, nonsignificant.

We also collected the mouse brain cortex tissue at three days and 28 days after skull marrow microinjections of spike S1, HA protein, and saline control to check for cell death and neuronal damage with the expression of cleaved caspase 3 and the Amyloid precursor protein (APP) respectively. Cleaved caspase 3 was increased in the brain three days after skull injection of both spike S1 and HA and remained elevated in the brain 28 days after spike S1 injection but returned to baseline after HA injection (**Fig. 5D-5F**). The spike S1 induced a significantly increased APP expression in the brain both three days and 28 days after injection, while the HA did not lead to any changes (**Fig. 5G-5I**).

These data demonstrated that the spike S1 protein alone locally residing in the skull marrow could signal into the brain to induce long-lasting pathological effects, including cell death and neuronal injury.

### Spike protein persists in the human skull marrow

Observing spike-induced changes in mice long after injection into the skull led us to test if we could also identify spike protein in the human skull long after infection. In Germany, 46% of the population reported having COVID-19^85^. Therefore, we hypothesized that if we sampled skull samples from several people who died from non-COVID reasons, we might detect the spike protein lingering in the skull marrow niches, which might be associated with ongoing neurological problems in COVID-19 patients.

We investigated the presence of spike protein in the skulls of 34 patients who died from non-COVID-related causes during the pandemic in 2021 and 2022. We indeed identified the spike protein in 10 of them (**Fig. 5J, Fig. S5A**). This refers to approximately 29% of people who potentially had COVID-19 in the past (without recognizing themselves or with no anamnestic data), and still having spike protein for the rest of their lives. These findings suggest that the spike protein persisted in the human skull beyond the viral detection time and recovery might be involved in long-term COVID-19 symptoms.

## DISCUSSION

The long-term complications of COVID-19 remain a major concern, and the multitude of symptoms reported suggest that these arise from effects of the infection that are only indirectly linked to the primary viral replication sites in the respiratory tract. For example, several neurological symptoms have been reported, most notably the long-lasting brain fog reported by many patients and significant brain tissue loss, even in mild COVID-19 cases, which urge exploration of the mechanisms of SARS-CoV-2 induced brain damage^86,87,8^.

Here, we used whole mouse tissue clearing technology to visualized the spike protein distribution by S1 protein with N501Y mutation, which has been shown to infect wild-type mice through binding with mouse ACE2^41,42^, to identify all potential tissue targets of the S1 protein and assess if S1 protein alone can induce brain pathologies in the absence of other viral proteins. In addition to the expected target organs such as the lung, kidney, and liver^88–90^, our results showed an accumulation of spike protein along with the viral nucleocapsid protein in the human skull marrow niches, SMCs, and meninges, suggesting that the virus might infect these tissues. However, additional data will be needed to show active replication. In contrast, only the spike protein was detected in the brain parenchyma. The detected spike protein could either be a residue of a viral infection of the brain that has been cleared or has infiltrated the brain from the cerebral circulation, in either case suggesting that the spike protein could have a long lifetime in the body^91^. This notion is supported by the observation that spike protein can be detected on patient immune cells more than a year after the infection^30^, and a recent preprint in medRxiv suggests spike protein’s persistence in plasma samples up to 12 months post-diagnosis^31^. The long-lasting spike protein near the brain tissue may explain some aspects of COVID-19 symptoms^31,92^. In our study, the persistence of spike protein in the skull marrow and brain tissue was identified also in post-mortem human samples. We also confirmed this phenomenon in K18-hACE2 mice infected with two different SARS-CoV-2 strains. It is worth noting that intravenous injection of spike protein induced a broad spectrum of proteome changes in the mouse skull marrow, meninges, and brain, including proteins related to coronavirus disease, complement and coagulation cascades, neutrophil degranulation, NETs formation, and PI3K-AKT signaling pathway, demonstrating the immunogenicity of SARS-CoV-2 spike protein in the absence of other viral components.

Although many studies in humans, animals, and cell line models addressed the molecular underpinning of SARS-CoV-2 infection, very few currently link the brain-associated pathologies evident in severe COVID-19 cases with the changes in the host proteome of the brain and adjacent tissues^93^. In this study, the spatial proteomics datasets of COVID-19-infected brain samples provide leads to study spike-specific changes in the brain. Our study benefited from the simultaneous analysis of compartments adjacent to COVID-19-infected individuals, namely the skull marrow and the meninges. The 29 proteins that the SARS-CoV-2 viral genome encodes^94^ directly or indirectly regulate the expression of many host proteins. Our molecular analysis suggests activation of immune response in the skull-meninges-brain axis, potentially via recruiting and increasing the activity of neutrophils similar to what has been reported in the respiratory tract^95,96^. Individual viral proteins have been suggested to exert physiological effects in the absence of other viral components, especially the spike protein which has been reported to induce the expression of inflammatory cytokines and chemokines in macrophages and lung epithelial cells and to compromise endothelial function^24–29^.

While investigating the changes in host cell expression, we find that specific host processes are most consistently dysregulated in individual tissues. In the skull marrow, predominant dysregulated pathways were the coronavirus disease pathway and complement and coagulation cascades, as reported previously for peripheral organs of patients with severe COVID-19^97^. The increased expression of pro-inflammatory proteins, including calprotectin, and proteins associated with thromboses, such as PF4 and PPBP, illustrate that the viral proteins act as an inflammatory stimulus that leads to the development of a significant immune response in the brain. However, we cannot distinguish between the direct effects of viral factors and the systemic effects of the disease. In the meninges, a significant consequence of the inflammatory state is an upregulation of proteins involved in neutrophil degranulation, such as PDAP1, DBNL, CD44, RAB6A, and HMGB1 that may be linked with a process known as NETosis to contain the infection. The upregulation of NET proteins presumably leads to high levels of NETs in the skull-meninges-brain axis and the infiltration of circulating neutrophils into the meninges. NETs could further propagate the inflammation^68^, potentially inducing tissue damage, including endothelium damage^98^, leading to pathologies such as thrombosis and alterations in the coagulation process^99^. Proteins related to the neurodegeneration pathway and damage to the blood-brain barrier were the most prominent dysregulated molecules in the brain.

The dysregulation of the complement and coagulation pathways was detected in both the skull marrow and the brain. This might explain the observed propensity of COVID-19 patients to develop mini-infarcts in the brain parenchyma^100,101^ and our observation of an increased level of micro-bleedings in COVID-19 patients, potentially contributing to the observed brain damage in the COVID-19 patients in acute or chronic stages.

All three tissues enriched proteins related to neutrophil degranulation and neutrophil extracellular trap formation. NETs formation in the skull marrow and neutrophil degranulation in the meninges suggest neutrophils may play a key role in maintaining inflammatory responses in and around the central nervous system. Reportedly, the viral spike protein leads to the activation of RHOA, which triggers the disruption of blood–brain barrier^102^. The RHOA GTPase identified in meninges is also reported to be regulated by PI3K^103^ and active in the neutrophil degranulation pathway to regulate actin dynamics and cell migration^103^. HSP90AA1 is another protein related to the PI3K-AKT signaling pathway previously associated with SARS-CoV-2 gene expression and disease severity^104^. HSP90AA1 links the PI3K-AKT signaling pathway to proteasome-mediated protein degradation, a pathway whose activation in COVID-19 infection is well established^105^. Other members of the heat shock protein family are overexpressed in subsets of tissues (HSPA8 and HSP90AB1). The transcription factor STAT1 is another common upregulated protein in the skull marrow and brain related to interferon-alpha/beta signaling. Reportedly, SARS-CoV-2 blocks the translocation of STAT1 to the nucleus to dampen the transcription of interferon response-related genes^106^. Another protein related to the interferon response was MX1, and it was identified in the skull marrow and brain samples. This protein has been reported to have antiviral functions^74^. Several proteins related to the IL-18 signaling were also identified in the skull marrow, including COL3A1, related to lung fibrosis in COVID-19 patients^107^ and ARF6, which has been suggested to mediate viral entry by regulating endocytosis^108^. Extracellular matrix proteins COL4A1 and COL4A2 were downregulated in the skull marrow. COL4A1 protein has a role in regulating cerebrovascular homeostasis^109^, although a role in COVID-19 pathogenesis has not been reported yet. In contrast to our findings at the protein level, at the transcript level, COL4A2 was reportedly downregulated in COVID-19 brain samples^110^, but no role in COVID-19 pathogenesis has been established.

We identified several candidate proteins with no previous association with COVID-19, especially those earlier associated with neurological diseases, such as Ras-related protein Rab-8B (RAB8B), Ras-related protein Rab-6 (RAB6A, RAB6B) and EF-hand domain-containing protein D2 (EFHD2). Notably, their role has been associated with disorders such as Parkinson’s disease, Alzheimer’s disease, and dementia^111,112^.

The proteins and pathways identified to be differentially regulated in the brain, skull marrow, and meninges provide leads to investigate the molecular mechanisms of immediate and long-term consequences of SARS-CoV-2 infections for the human brain. The common dysregulation of coronavirus disease, complement activation, and PI3K-AKT pathways in the skull, meninges, and brain tissue demonstrated a common effect of SARS-CoV-2 infections on the immune system along the skull-meninges-brain axis. These molecules or molecular pathways can be leveraged as therapeutic targets to prevent or treat brain-related complications in COVID-19.

Our data may also suggest a mechanism for the virus’s entry into the central nervous system. In both mouse and COVID-19 human tissues, we find spike protein in the SMCs, which the virus or virus components could use to travel from the skull marrow to the meninges and the brain parenchyma^35–38^. Of course, the virus might take other routes to reach the brain in a not mutually exclusive way. For example, the virus could traverse the cerebrovasculature to reach the brain parenchyma or be carried there by immune cells (via neutrophils or phagocytic cells). More data will be needed to establish the most common route of brain invasion by SARS-CoV-2, which might differ between different parts of the brain. Brain invasion of virus-shed spike protein found in some COVID-19 cases has been linked to a compromised blood-brain barrier^34,113^ and trafficking along the olfactory nerve or vagus nerve^9^. Here, we suggest an alternative scenario wherein SARS-CoV-2 spike protein reaches first the skull marrow and then the meninges before entering the brain.

Spike-induced alterations in the skull-meninges-brain axis present diagnostic and therapeutic opportunities as both skull and meninges are easier to access than brain parenchyma. Panels of such proteins tested in plasma samples of COVID-19 patients might provide an early prognosis of brain-related complications. Our future efforts will be committed to characterizing these proteins towards their use as biomarkers and therapeutic targets for neurological dysfunction in COVID-19 infections.

## MATERIALS AND METHODS

### Animals involved in the study

We used the following animals in this study: 2 months old wildtype C57Bl6/J mixed sex mice purchased from Charles River Laboratories (MA, USA). Animals were housed on a 12/12 hr light/dark cycle and had arbitrary access to food and water. Temperature was maintained at 18-23 °C and humidity at 40-60%. Animal experiments were performed according to institutional guidelines of the Helmholtz Center Munich and Ludwig Maximilian University of Munich after approval by the Ethical Review Board of the Government of Upper Bavaria (Regierung von Oberbayern, Munich, Germany) and according to the European Directive 2010/63/EU for animal experiments. All data are reported according to the criteria of ARRIVE. The sample size was chosen based on previous experience with similar models.

### Origin of human tissue and related work

PFA-fixed human brains and human skull blocks were obtained from donors and autopsies in case of a diagnosis of COVID-19 during a lifetime or positive SARS-CoV-2 qPCR test post-mortem and in accordance with the European Control for Infectious Diseases. All donors or next-of-kin gave their informed and written consent to examine the cadavers for research and educational purposes. Human brain tissue was obtained from the Anatomical Institute of the University of Leipzig, Germany, and the Institute of Pathology of the Technical University of Munich, Germany. Skull samples were collected during autopsies at the Institute of Legal Medicine of the University Medical Center Hamburg-Eppendorf, Germany. The post-mortem interval was on average 5 d. Institutional approval was obtained in accordance with the 1994 Saxon Death and Burial Act and of the independent ethics committee of the Hamburg Chamber of Physicians (protocol 2020-10353-BO-ff). Patient information can be found in the supplementary material (Table. 1).

### Spike S1 protein whole mouse trafficking

Spike S1 protein (N501Y), Spike S1 protein (WT), and hemagglutinin (HA) were labeled with the fluorescent dye Alexa Fluor 647 according to the manufacturer’s protocol (Alexa Fluor 647 conjugation kit lightning link, Abcam, ab269823). Briefly, the candidate protein (20 μg) supplied by the manufacturer (SARS-CoV-2 (COVID-19) S1 protein (N501Y), His Tag, Variant Lineage B.1.1.7 (S1N-C52Hg, Acrobiosystems), HA Recombinant Influenza A Virus Protein, subtype H1N1 (A/California/04/2009) (A42579, Life Technologies), Coronavirus (COVID-19 Spike Protein; Full Length) Antigen, Recombinant >90% (ECB-LA636-100, Enzo Life Sciences)) were dissolved in 0.1 M PBS (18 μl, pH 7.4). The modifying reagent (1 μl) was added to the protein solution. The solution was then mixed gently and transferred to a lyophilized material. After 15 minutes of incubation at room temperature, the reaction was terminated by adding the quencher reagent (1 μl). The fluorine-labeled protein had a concentration of 1 mg ml-1. Anesthesia was induced in the animals with 4% isoflurane in an N2O/O2 mixture (70%/30%) and then maintained with 1.5% isoflurane in the same mixture for the entire injection. A total of 0.1 ml of PBS solution containing 1 μg Alexa-647 conjugated protein was injected intravenously into the mouse tail vein. The protein was allowed to circulate throughout the mouse for 30 minutes. Mice were then transcardially perfused as described in the perfusion and tissue preparation section below.

### Intravenous injection and skull marrow microinjection

Intravenous injections were performed through the tail vein under general anesthesia using a 1 mL syringe with a 28-gauge needle. Spike S1 protein (N501Y) and hemagglutinin (HA) were dissolved in saline to a stock concentration of 1 mg ml-1, a total of 100 μl saline containing 10 μg protein was injected to each mouse. The mice were perfused 3 days post injection, skull, meninges, and brain were then collected for proteomic analysis.

Microinjection into the skull marrow was performed in mice under 1.5% isoflurane anesthesia using a 5 μl syringe (#87943, Hamilton) equipped with a 34-gauge blunt needle. To expose the anterior marrow sites (near the bregma), the skin above the skull was opened at the midline. A drill bit was used as a pilot drill to remove the cortical portion of the skull and expose the spongy bone marrow. A microinjection was performed for each marrow site pre-drilled with a 30-gauge needle (#305106, BD) to prevent perforation of the inner skull wall. In this study, the red tracker (C34565, Molecular Probes) was diluted to 10 μM according to the manufacturer’s protocol. The Alexa Fluor 647 conjugated protein solution (1 mg ml-1) was prepared and a total volume of 2.5 μl solution was slowly injected per injection site through the pre-drilled sites (20-30 s per injection), resulting in a total of 10 μl of conjugated spike protein injected per skull. The injection process was closely monitored under a microscope. Animals in which the procedure was unsuccessful at this time were removed from the study. After the injection was completed, the skin was sutured with 6-0 silk. The mice were euthanized at given time points and the heart was then perfused with PBS and fixed with 4% PFA for further analysis.

### Perfusion and tissue preparation

Mice were deeply anesthetized by intraperitoneal injection with a combination of midazolam, medetomidine, and fentanyl (1 ml/100 g of body mass for mice) until the mice showed no response with a pedal reflex. Then, the mice were perfused intracardially with heparinized 0.1 M PBS (10 U/ml of Heparin, Ratiopharm, N68542.03; 100-125 mmHg pressure using a Leica Perfusion One system) for 5 minutes to remove the blood, followed by a total volume of 20 ml of dextran (5mg/ml, MW 500000, Sigma, 52194) for vascular labeling. Mice were then perfused with 50 ml of 4% paraformaldehyde (PFA) in 0.1 M PBS (pH 7.4; Morphisto, 11762.01000) for fixation. Then, bodies were post-fixed in 4% PFA for 1 day at 4°C and later washed three times with 0.1 M PBS for 10 minutes at room temperature. The mice were then tissue cleared using the 3DISCO method.

### Clearing the whole mouse using 3DISCO method

We performed tissue clearing based on the 3DISCO protocol for whole mice as previously described. For this purpose, mouse bodies were placed in a 300 ml glass chamber and incubated in 200 ml of the following gradient of THF (Tetrahydrofuran, Roth, CP82.1) in distilled water in a fume hood with gentle shaking: 50% ×1, 70% ×1, 80% ×1, 100% ×2, 12 h for each step, followed by 3 h in dichloromethane (DCM, Sigma, 270997) and finally in BABB solution (benzyl alcohol + benzyl benzoate 1:2, v/v, Sigma, 24122 and W213802) until optical transparency.

### Enzyme-Linked ImmunoSorbent Assay (ELISA)

Mouse blood was collected into citrate-treated tubes and centrifuged at 2000 x g for 15 minutes, the supernatant was immediately proceeded for ELISA assay according to the manufacturer’s protocol. Mouse Interferon Gamma (IFNg) ELISA Kit (RD-IFNg-Mu, Reddot biotech), Mouse Interleukin 6 (IL6) ELISA Kit (RD-IL6-Mu, Reddot biotech), Mouse Interferon Gamma Induced Protein 10kDa (IP10) ELISA Kit (RD-IP10-Mu, Reddot biotech) were used in this study.

### Deep tissue immunolabelling of human tissue

Deep tissue labeling was performed according to the protocol known in our laboratory as the SHANEL protocol. Cut 1 cm thick slices of the desired brain tissue or human skull before SHANEL pretreatment. Then treat the human brain slices with CHAPS /NMDEA solution twice at room temperature for 12 hours each time. Wash three times with PBS for 20 minutes each time. Tissue was dehydrated by stepwise addition of ethanol (50% ×1, 70% ×1, 100% ×1, 4 h for each step). Change to DCM/MeOH (2:1, v/v) solution overnight. Tissues were rehydrated stepwise in diH2O to remove ethanol (100% ×1, 70% ×1, 50%×1, 4 h for each step). Treat the tissue overnight with 0.5 M acetic acid solution, wash twice for 20 min in diH2O. Treat the tissue with guanidine solution for 6 h, and wash 2 more times in diH2O. Block the tissue with blocking buffer (0.2% Triton X-100, 10% DMSO (Roth, A994.2), 10% goat serum in 0.1 M PBS) at 37 °C overnight and then incubate it with propidium iodide (1:1000, Sigma, P4864) or antibodies against SARS-CoV-2 (COVID-19) Spike (1:1000, GeneTex, GTX135356, GTX632604), CD31 (1:1000, Abcam, ab32457), Nucleocapsid (1:1000, Invitrogen, PA1-41098), ACE2 (1:1000, Invitrogen, PA5-20039), CD45 (1:1000, 14-0451-85, ThermoFisher), Iba1 (1:1000, 019-19741, Wako) in antibody incubation buffer (3% goat serum, 3% DMSO, 0.2% Tween-20, 10 mg L-1 heparin in 0.1 M PBS) at 37 °C for 5 days. After incubation with the primary antibodies, the tissue was washed twice in PBS for 1 hour, and the secondary antibodies (goat anti-rabbit IgG Alexa Fluor 647, Invitrogen, A-21245; goat anti-rabbit IgG Alexa Fluor 568, Invitrogen, A-11036; goat anti-mouse IgG Alexa Fluor 568, Invitrogen, A-11031; goat anti-mouse IgG Alexa Fluor 647, Invitrogen, A-21235; goat anti-rat IgG Alexa Fluor 568, Invitrogen, A-11077) were incubated at a concentration equal to that of each primary antibody for 5 days at 37 °C. Wash the samples three times for 1 hour at room temperature with wash buffer to remove excess antibody. Dehydrate gradually in EtOH/H2O (50% ×1, 70% ×1, 80% ×1, 100% ×2, 6 h for each step). Switch to DCM for 1 h and then to BABB until the sections become transparent. Finally, examine the results with a light-sheet microscope for volume imaging.

### Light-sheet microscopy imaging

Image stacks were acquired using a Blaze ultramicroscope or II ultramicroscope (LaVision BioTec) with an axial resolution of 4 μm and the following filter sets: ex 470/40 nm, em 535/50 nm; ex 545/25 nm, em 605/70 nm; ex 640/40 nm, em 690/50 nm. Entire mouse bodies were scanned individually with a low magnification Ultramicroscope Blaze light sheet microscopy objective: 1.1× objective (LaVision BioTec MI PLAN 1.1×/0.1 NA [WD = 17 mm]). We covered the entire mouse with 3 × 8 tile scans with 25% overlap and imaged from the ventral and dorsal surfaces to 11 mm in depth, covering the entire body volume with a Z-step of 10 μm. The width of the light-sheet was kept at 100% and the exposure time was set at 120 ms. The laser power was adjusted depending on the intensity of the fluorescence signal to avoid saturation. High-magnification tile scans for multiple organs (including brain, heart, lung, thymus, liver, spinal cord, spleen, kidney, intestine, testis, and ovary) were acquired individually with high-magnification objectives (Olympus XLFLUOR 4× corrected/0.28 NA [WD = 10 mm] and PLAN 12×/0.53 NA [WD = 10 mm], LaVision BioTec MI) coupled to an Olympus rotary zoom unit (U-TVCAC) set at 1×. The high magnification tile scans were acquired with a 20% overlap and the width of the light-sheet was reduced to achieve maximum illumination in the field of view. The acquired raw images TIFF were processed with Fiji’s stitching plugin, then with Vision4D (v.3.3 × 64, Arivis) for volume fusion, and visualized in Imaris (v.9.6 × 64, Imaris) for 3D reconstruction, analysis, and video generation.

### Immunofluorescence and confocal microscopy

Briefly, frozen sections were treated with 0.2% Triton X-100 in PBS for 15 minutes and blocked with 1 serum in PBST for 40 minutes at room temperature. They were then incubated with primary antibodies, SARS-CoV-2 (COVID-19) Spike (1:500, GeneTex, GTX135356, GTX632604), Nucleocapsid (1:500, Invitrogen, PA1-41098), ACE2 (1:500, Invitrogen, PA5-20039), NeuN (1:500, Invitrogen, PA5-78499), Anti-F4/80 antibody [CI:A3-1] - Macrophage Marker (ab6640, Abcam), Anti-Influenza A H1N1 hemagglutinin antibody (ab128412, Abcam), Recombinant Anti-Sodium Potassium ATPase antibody[EP1845Y] - Plasma Membrane Loading Control (ab76020, Abcam). overnight at 4°C and washed with PBS for 15 min, Alexa-conjugated secondary antibodies (1:1000, goat anti-rabbit IgG Alexa Fluor 647, Invitrogen, A21245; goat anti-rabbit IgG Alexa Fluor 568, Invitrogen, A-11036; goat anti-mouse IgG Alexa Fluor 568, Invitrogen, A-11031; goat anti-mouse IgG Alexa Fluor 647, Invitrogen, A-21235; goat anti-rat IgG Alexa Fluor 568, Invitrogen, A-11077) were incubated for 1 hour at room temperature. Sections were mounted after staining with Hoechst 33342 (Invitrogen). Images were acquired with a 40x immersion objective (Zeiss, EC Plan-Neofluar 40x/1.30 Oil DIC M27) of a confocal microscope (ZEISS LSM880).

To detect spike protein in each sample, three random regions were selected, and 20 sections were prepared for each region.

### Immunohistochemistry

Briefly, frozen sections were permeabilized in 0.1% Triton X-100 in PBS, treated with 3% H2O2 for 10 min, and blocked with 10% goat serum for 20 min, incubated with primary antibody against SARS-CoV-2 (COVID-19) Spike (1:200, GeneTex, GTX135356) at room temperature for 1 h. Staining was detected with goat anti-rabbit IgG HRP antibody (1:200, Abcam, ab6721) and revealed by incubation with diaminobenzidine for 10-20 seconds (Vector, VEC-MP-7714). Hematoxylin (Sigma, 51275) was used as counterstaining. Prussian blue staining was performed according to manufacturer’s instruction (NovaUltra, IW-3010).

### SARS-CoV-2 real-ti transcription

#### polymerase chain reaction (qRT -PCR) test

Decalcified COVID-19 skull samples were minced in PBS, COVID-19 meninges samples were ground in liquid nitrogen and dissolved in PBS, the tissue extract was passed through 40 μm strainer. RNA extraction was performed using RNeasy FFPE Kit (QIAGEN, 73504), followed by SARS-CoV-2 qRT-PCR using the Seegene Allplex™ 2019-nCoV Assay (cat. no: RP10243X) on a CFX96 Real-time PCR Detection System-IVD (Bio-Rad)

### Laser capture microdissection for spike positive and negative regions

For microdissection of the spike protein-positive regions, we laser-cut and isolated the selected samples using the laser capture system (Leica, LMD7000). Briefly, the cryosections of human brain were mounted on a Polyethylene naphthalate (PEN) Membrane Slide (Zeiss, 415190-9041-000) and then stained with spike protein antibodies using the IHC kit (Abcam, ab210062). After further processing, the sections were serially dehydrated with ethanol and air-dried under a fume hood for 15 minutes. The spike protein-positive and spike protein-negative regions of the COVID-19 human brain sections were selected with a closed-shape manual drawing tool and dissected using a UV laser. The excised regions were collected into a 0.5 ml tube and examined by camera. An accumulated area of 6 mm2 was collected by laser cut and using 40× objective (HC PL FL L 40×/0.60 XT CORR). The tissues were quickly spun down and stored at -80 °C for further proteomic analysis.

## Statistical analysis

Prism 8.0 was used for all statistical calculations (GraphPad Software, San Diego, CA). Two-tailed *t* tests were used to compare two means. The statistical tests used are specified in all figure legends. The data were analyzed with the Shapiro-Wilk normality test and had Gaussian distribution.

### Sample preparation for mass spectrometry analysis

The human samples comprised post-mortem samples of human skull and meninges tissues from ten COVID-19 deceased and ten control donors. Protein extraction was carried out from 4% neutral buffered formalin-fixed tissues. Briefly, the cells from the skull and the meninges tissues were isolated by mincing or grinding, and the tissue extract was passed through a 40 μm strainer. The cell pellet was washed with PBS and resuspended in SDS-lysis buffer (6% Sodium dodecyl sulfate, 500 mM TrisHCl, pH 8.5). This was followed by heating at 95°C for 45 min at 1000 rpm in a thermomixer. The samples were then subjected to ultrasonication using a Bioruptor Pico sonication device operated at high frequency for 30 sec on and off for 30 cycles. After ultrasonication, the samples were again heated at 95°C for 45 min at 1000 rpm in a thermomixer. This was followed by protein precipitation in ice-cold acetone (80% v/v) overnight at -80°C, followed by centrifugation for 15 min at 4°C. For reduction and alkylation, the proteins were resuspended in SDC buffer and heated at 95°C for 10 min with 1000 rpm. Trypsin and LysC digestion were carried out at an enzyme/substrate ratio of 1:50 and the samples were incubated at 37°C overnight, 1000 rpm in a thermomixer. Next, peptides were acidified using 1% TFA in 99% isopropanol in a 1:1 v/v ratio. The peptides were subjected to in-house built StageTips consisting of two layers of styrene-divinylbenzene reversed-phase sulfonate (SDB-RPS; 3 M Empore) membranes. Peptides were loaded on the activated (100% ACN, 1% TFA in 30% Methanol, 0.2% TFA, respectively) StageTips, run through the SDB-RPS membranes, and washed by EtOAc including 1% TFA, isopropanol including 1% TFA, and 0.2% TFA, respectively. Peptide elution was carried out in 60 μL of 1.25% Ammonia, 80% ACN and dried for 40 min at 45 °C in a SpeedVac (Eppendorf, Concentrator plus). The dried peptides were reconstituted in 10 μL of 2% ACN/0.1% TFA and peptide concentration was estimated using Pierce™ Quantitative Colorimetric Peptide Assay.

### Liquid chromatography and mass spectrometry (LC-MS/MS)

The mass spectrometry data was generated in both data-dependent acquisition (DDA) as well as data-independent acquisition (DIA) modes. For DDA analysis, EASY-nLC 1200 (Thermo Fisher Scientific) combined with an Orbitrap Exploris 480 Mass Spectrometer (Thermo Fisher Scientific) and a nano-electrospray ion source (Thermo Fisher Scientific) was used. Peptides were separated by reversed-phase chromatography using a binary buffer system consisting of 0.1% formic acid (buffer A) and 80% ACN in 0.1% formic acid (buffer B) with a 120 min gradient (5-30% buffer B over 95 min, 30-65% buffer B over 5 min, 65-95% buffer B over 5 min, wash with 95% buffer B for 5 min) at a flow rate of 300 nL/min.

MS data were acquired using a data-dependent cycle time (1 second) scan method. Full scan MS targets were in a 300–1650 m/z scan range with a normalized automatic gain control (AGC) target %300. The data was acquired at 60000 resolution with a 25 ms maximum injection time. Precursor ions for MS/MS scans were fragmented by higher-energy C-trap dissociation (HCD) with a normalized collision energy of 30%. MS/MS scan sets were 15000 resolution with an AGC target %100 and a maximum injection time of 28 ms. For DIA analysis, the LC-MS/MS analysis was carried out using EASY nanoLC 1200 (Thermo Fisher Scientific) coupled with trapped ion mobility spectrometry quadrupole time-of-flight single cell proteomics mass spectrometer (timsTOF SCP, Bruker Daltonik GmbH, Germany) via a CaptiveSpray nano-electrospray ion source. Peptides (50 ng) were loaded onto a 25 cm Aurora Series UHPLC column with CaptiveSpray insert (75 μm ID, 1.6 μm C18) at 50°C and separated using a 50 min gradient (5-20% buffer B in 30 min, 20-29% buffer B in 9 min, 29-45% in 6 min, 45-95% in 5 min, wash with 95% buffer B for 5 min, 95-5% buffer B in 5 min) at a flow rate of 300 nL/min. Buffer A and B were water with 0.1 vol% formic acid and 80:20:0.1 vol% ACN:water:formic acid, respectively. MS data were acquired in single-shot library-free DIA mode and the timsTOF SCP was operated in DIA/parallel accumulation serial fragmentation (PASEF) using the high sensitivity detection-low sample amount mode. The ion accumulation and ramp time was set to 100 ms each to achieve nearly 100% duty cycle. The collision energy was ramped linearly as a function of the mobility from 59 eV at 1/K0 = 1.6 V-s cm−2 to 20 eV at 1/K0 = 0.6 V-s cm−2. The isolation windows were defined as 24 × 25 Th from m/z 400 to 1000.

### Proteomics data processing

DDA files were processed with MaxQuant version 1.6.14.0. Default settings were kept if not stated otherwise. FDR 0.01 was used for filtering at protein, peptide, and modification levels. As variable modifications, acetylation (protein N-term) and oxidized methionine (M), as fixed modifications, carbamidomethyl (C) were selected. Trypsin/P and LysC proteolytic cleavages were added. Missed cleavages allowed for protein analysis was

2. “Match between runs” and label free quantitation (LFQ) were enabled, and all searches were performed against the human and mouse Uniprot database. diaPASEF raw files were searched against the human and mouse Uniprot database using DIA-NN^114^. Peptide length range from seven amino acids was considered for the search including N-terminal acetylation. Oxidation of methionine was set as variable modifications and cysteine carbamidomethylation as fixed modification. Enzyme specificity was set to Trypsin/P with 2 missed cleavages. The FASTA digest for the library-free search was enabled for predicting the library generation. The FDR was set to 1% at precursor and global protein levels. Match-between-runs (MBR) feature was enabled and the quantification mode was set to “Robust LC (high precision)”. The Protein Group column in DIA-NN’s report was used to identify the protein group and PG.MaxLFQ was used to calculate the differential expression.

### Proteomics downstream data analysis

Proteomics data analysis was performed in Perseus and R studio. The protein groups were filtered such that only those proteins were considered for differential expression which was present in 3 out of 4 samples in each group with valid values. The values were log2 transformed and normalized with median centering the dataset. The missing values were randomly drawn from a Gaussian distribution with a width of 0.3 standard deviations that was downshifted by 1.8 standard deviations. The correlation heatmap was computed using the Pearson correlation coefficient. Differential protein expression between the COVID-19 and control groups was performed by student-t test. The Benjamini-Hochberg procedure was applied to correct for multiple comparisons. p-value < 0.05 and fold change > 1.5 was taken as statistical significance, unless specified. The gene ontology (GO) and pathway enrichment were carried out in Cytoscape using ClueGO plug-in^115^ or ClusterProfiler. PCA plot, volcano plot, GO_ Chord plot, and heatmap were generated in R studio.

## ACKNOWLEDGEMENTS

We thank Alireza Ghasemi for developing the python script to stitch sequences of images. We thank Christoph Krisp (Bruker Daltonics) for helpful discussions on mass spectrometer data acquisition. We thank Markus Elsner for revising the manuscript. Illustrations were created with BioRender.com. Z.R. and H.M. would like to thank the China Scholarship Council (CSC) for the financial support (No. 201806310110 and No. 201806780034). We acknowledge all families supporting our research after losing their beloved ones during the pandemic.

This work was supported by the Vascular Dementia Research Foundation, Deutsche Forschungsgemeinschaft (DFG, German Research Foundation) under Germany’s Excellence Strategy within the framework of the Munich Cluster for Systems Neurology (EXC 2145 SyNergy, ID 390857198) and DFG (SFB 1052, project A9; TR 296 project 03) as well as German Federal Ministry of Education and Research (Bundesministerium für Bildung und Forschung, BMBF) within the NATON collaboration (01KX2121).

## AUTHOR CONTRIBUTION

A.E. conceived and led all aspects of the project.

Z.R. and H.M. designed and carried out most of the experiments. S.K., S.U., and Ö.S.C. performed mass spectrometry-based proteomics experiment. Z.R., H.M., and S.K. performed data analysis and visualization. V.G.P., J.S., J.V., T.B.H., and B.O. helped and organized human sample collection.

J.C. and J.V. dissected the human skull with dura mater sample and collected clinical information. C.D., H.S., H.F., K.S., and I.B. provided human brain and skull samples. J.M.W. performed RT-PCR analysis. F.C. assisted in sample homogenization.

Z.I.K. collected and processed human skull samples. M.A., I.H., and S.Z. participated in prototyping experiments. A.E., F.H., and H.S.B. supervised the project. Z.R., H.M., and S.K. wrote the first draft of manuscript. A.E. wrote the final manuscript. All authors reviewed and approved the final manuscript.

## DATA AVAILABILITY

The mass spectrometry raw files and the search output files are available at http://www.ebi.ac.uk/pride/archive/ with the PRIDE dataset identifier PXD034178.

## VIDEO LEGENDS

**Video 1**. High resolution visualization of spike S1 protein in mouse organs obtained by light-sheet microscopy. Section view in liver, intestine, heart, spleen, pancreas. The spike S1 protein is shown in green and dextran vessel labeling in magenta.

**Video 2**. 3D animation of mouse reproductive system scanned by light-sheet microscopy. The background is shown in gray, the spike S1 protein are shown in green and dextran vessel labeling in magenta.

**Video 3**. 3D reconstruction of an intact mouse torso scanned by light-sheet microscopy. The background is represented in gray, the spike S1 protein in yellow glowing color. Spike S1 protein in various organs are visualized in high contrast over the background.

**Video 4**. Sagittal view inspection of the cleared mouse head reveals the spike S1 protein in the mouse skull marrow, skull-meninges-connections and meninges. The background is shown in gray, the spike S1 protein are shown in green and dextran vessel labeling in magenta.

**Video 5**. 3D animation of mouse central nervous system scanned by light-sheet microscopy. The background is shown in gray, the spike S1 protein are shown in green and dextran vessel labeling in magenta.

**Video 6**. 3D reconstruction of COVID-19 patient skull with dura sample shows spike protein in human skull marrow, skull-meninges-connections and meninges. The background is shown in blue, the spike protein in red and PI labeling of nucleus in cyan, dura mater is shown in yellow.

**Video 7**. Section view of COVID-19 patient skull sample shows spike protein in the skull marrow. The background is shown in gray, the spike protein in red.

**Video 8**. 3D reconstruction of COVID-19 patient brain with meninges sample. Section view shows spike protein in meninges and brain vasculature. The background is shown in gray, the spike protein in red and Iba1 labeling of myeloid cells in cyan.

**Video 9**. 3D reconstruction of COVID-19 patient brain cortex shows spike protein in brain vasculature and parenchyma. The background is shown in gray, the spike protein in green.

**Figure S1.**
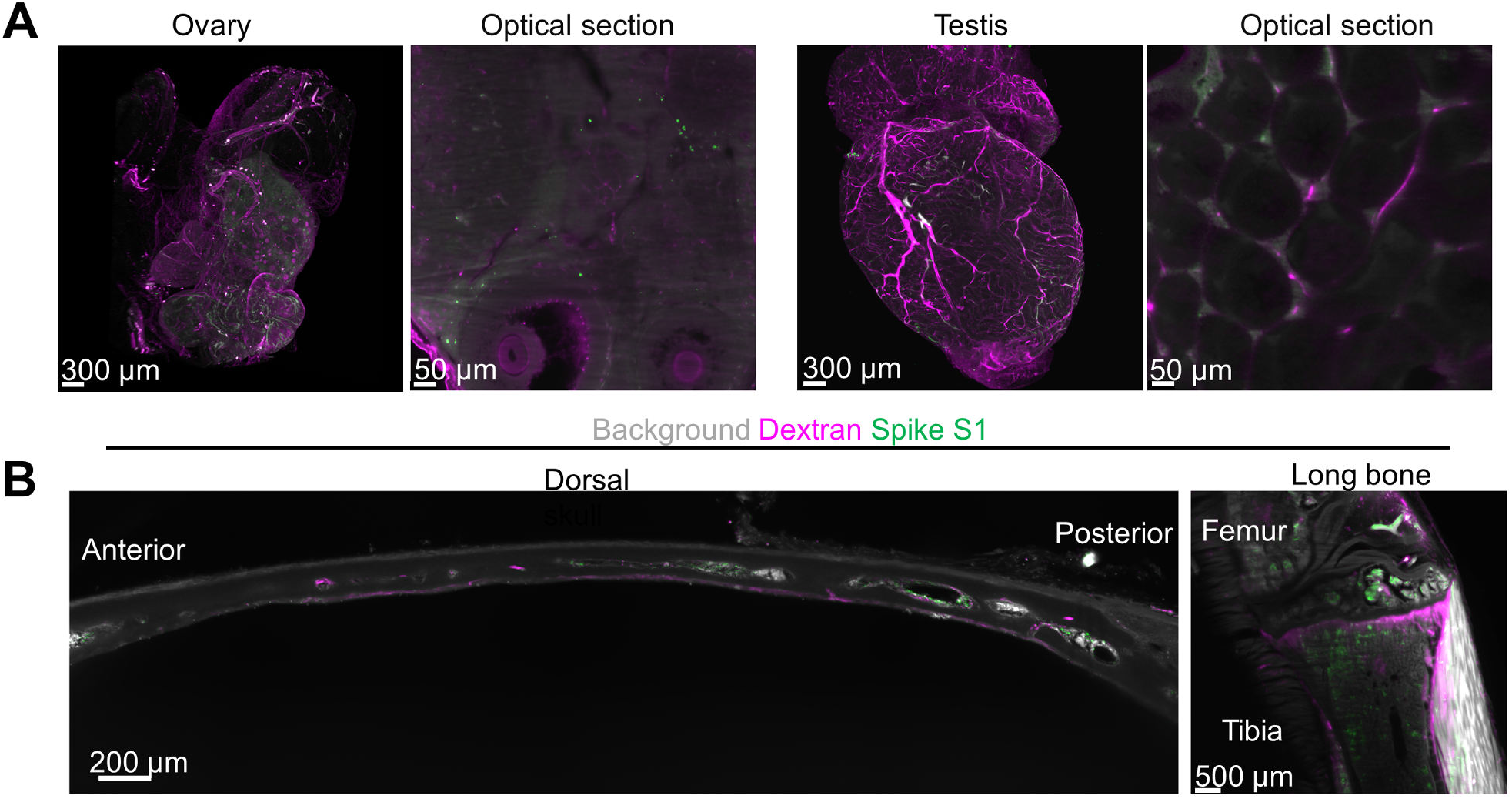
SARS-CoV-2 spike S1 protein distribution. (A) Spike S1 protein in mouse ovary and testis. (B) Representative images of spike S1 protein in mouse skull marrow and bone marrow of tibia and femur.

**Figure S2.**
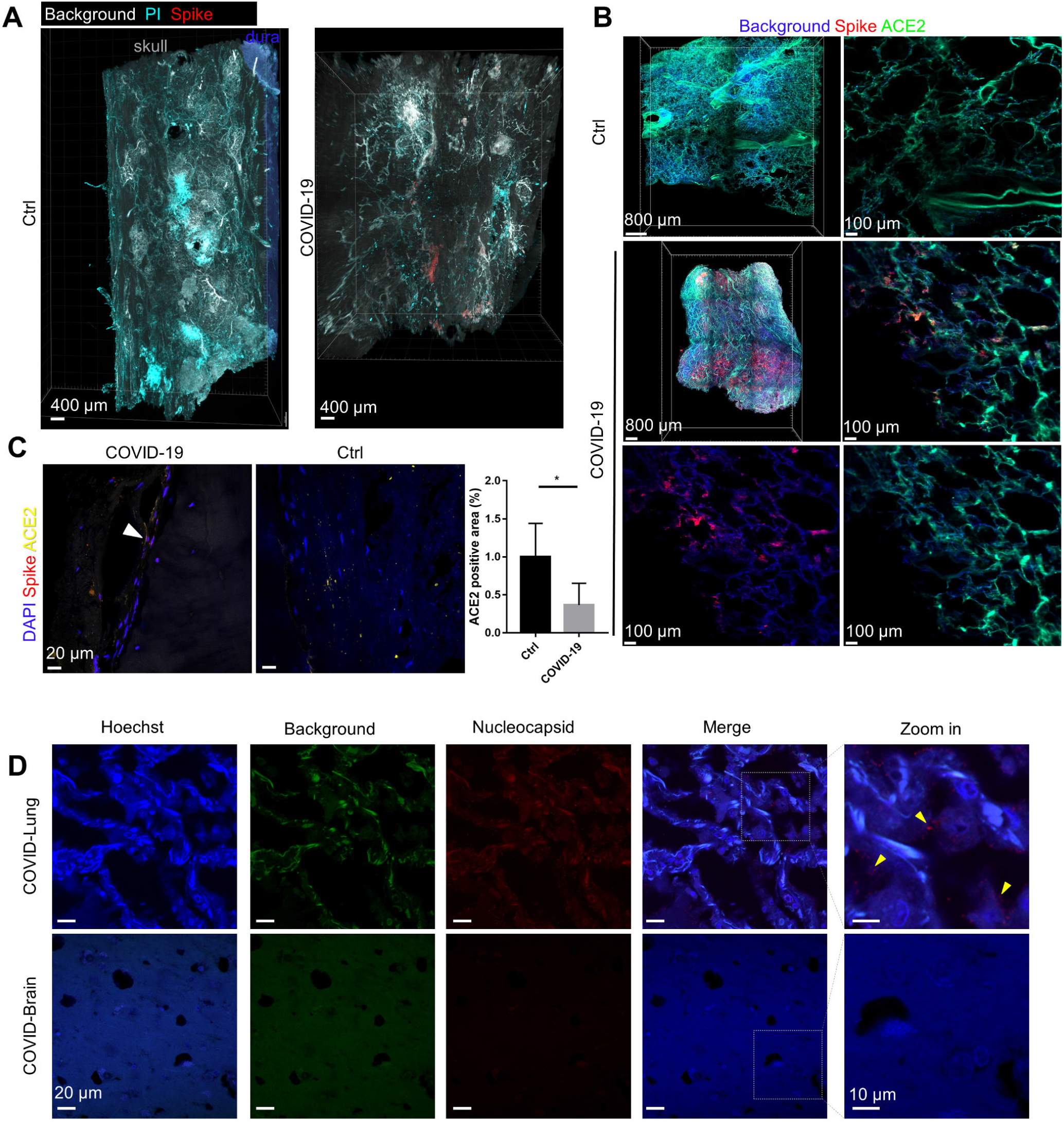
Additional microscopy images of COVID-19 patient skull, meninges, brain, and lung. (A) Spike protein staining in human skull samples from COVID-19 patient and non-COVID-19 control patient. (B) Representative images of spike protein and ACE2 staining in human lung tissue block. (C) Representative confocal images of spike protein antibody and ACE2 antibody labeling in the COVID-19 patient and control meninges. Quantification of the ACE2 positive area in 6 fields of view. n = 3. Data are mean ± SD. Two-tailed unpaired Student’s t test. *p < 0.05. (D) Representative confocal images of nucleocapsid staining in COVID-19 patient lung and brain tissue.

**Figure S3.**
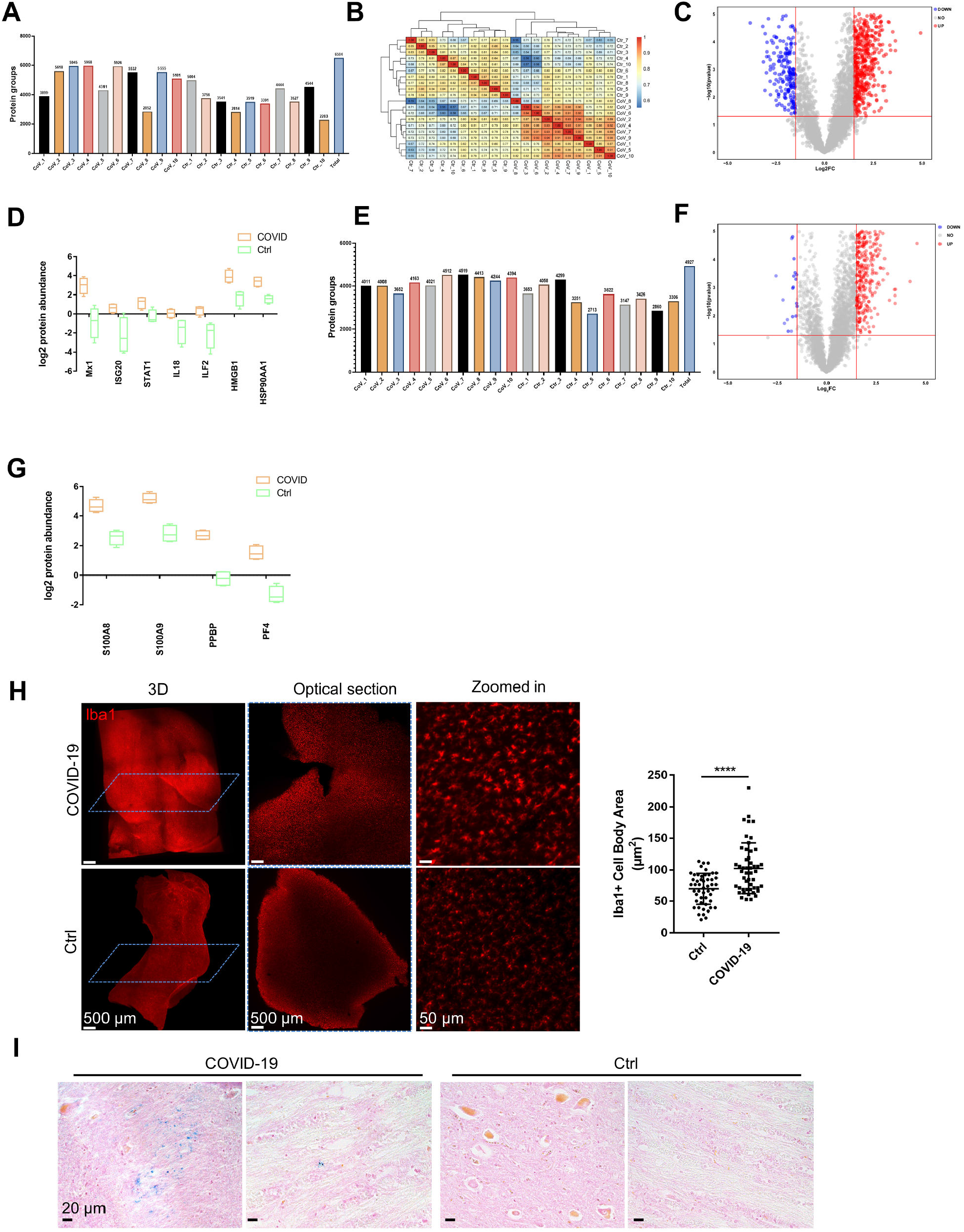
Additional information of mass-spec based proteomic and histology on human samples. (A) Protein groups identified in human skull marrow samples. (B) Pearson correlation analysis of skull marrow samples. (C) Volcano plot between the skull marrow samples of COVID-19 patients and controls. (D) Dysregulated proteins of interest in skull marrow samples. (E) Protein groups identified in human meninges samples. (F) Volcano plot between the meninges samples of COVID-19 patients and controls. (G) Dysregulated proteins of interest in meninges samples. (H) Representative images of Iba1 antibody staining and light-sheet imaging of human brain cortex. Non-COVID-19 sample is used as control. Quantification of microglia cell body area, each filled dot represents the area of an individual microglia soma. n = 3. Graph displays mean ± SEM. Two-tailed unpaired Student’s t test, ****p < 0.0001. (I) Representative images of Prussian blue staining in COVID-19 patient and control brain. Cell nucleus in red.

**Figure S4.**
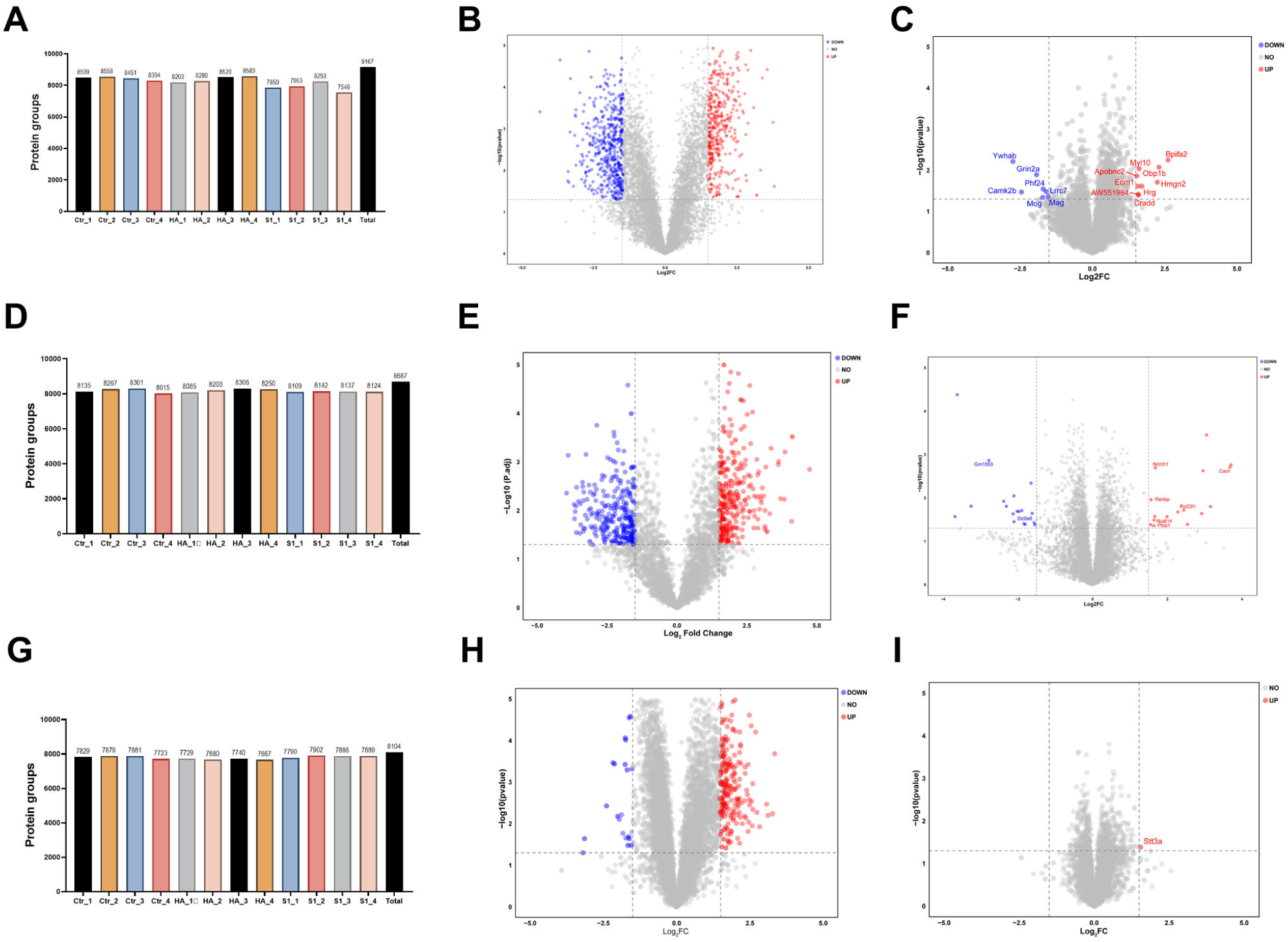
Information of Mass-spec based proteomic on mouse skull marrow, meninges, and brain. (A) Protein groups identified in the mouse skull marrow 3 days after i.v. injection of spike S1, HA, and saline control. (B) Volcano plot between skull marrow samples after spike S1 injection and control (p < 0.05, Log2FC = 1.5). (C) Volcano plot comparing skull marrow samples after HA injection and control (p < 0.05, Log2FC = 1.5). (D) Protein groups identified in the mouse meninges 3 days after i.v. injection of spike S1, HA, and saline control. (E) Volcano plot between meninges samples after spike S1 injection and control (p < 0.05, Log2FC = 1.5). (F) Volcano plot comparing meninges samples after HA injection and control (p < 0.05, Log2FC = 1.5). (G) Protein groups identified in the mouse brain 3 days after i.v. injection of spike S1, HA, and saline control. (H) Volcano plot between brain samples after spike S1 injection and control (p < 0.05, Log2FC = 1.5). (I) Volcano plot comparing brain samples after HA injection and control (p < 0.05, Log2FC = 1.5).

**Figure S5.**
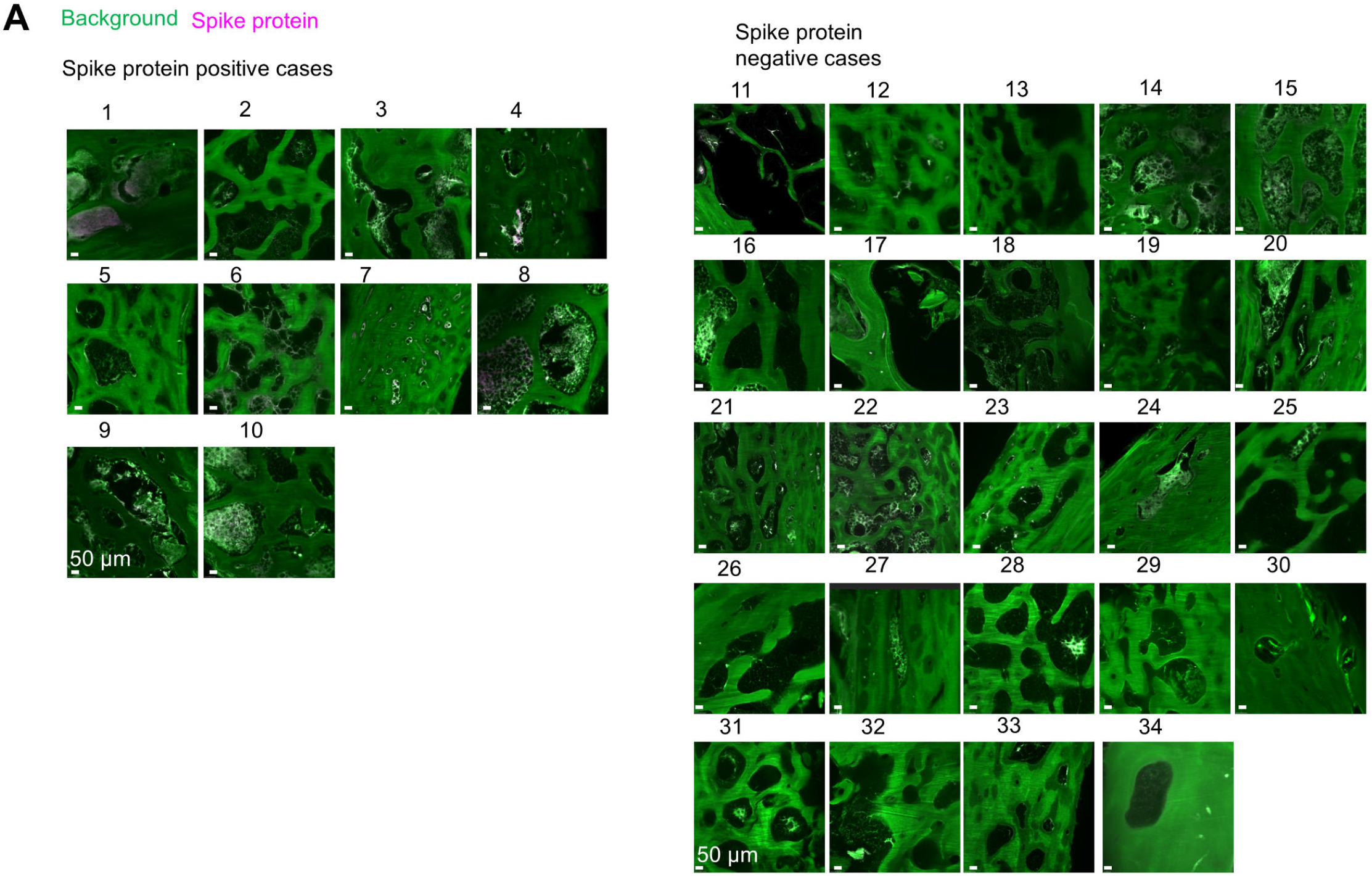
SARS-CoV-2 spike protein in the human skull. (A)Representative images of spike protein staining in skull samples from patients died during COVID-19 pandemic, where the cause of death was not COVID-19. A Z-stack of 1200 μm thick consecutive sections were inspected.

