## Supplementary material for "SARS-CoV-2 Spike Protein Accumulation in the Skull-Meninges-Brain Axis: Potential Implications for Long-Term Neurological Complications in post-COVID-19": Table 1

| Group | Number | Sex | Age range | PMI | Comorbidities | Brain weight[g] | Days of disease | PCR |
|---|---|---|---|---|---|---|---|---|
| COVID-19 skull/meninges | | | | | | | | |
| | 1 | f | 80~100 | 5 | Uterus myomatosus, Diverticulosis, Gall bladder stones | 1230 | n/a | skull-, meninges-, brain n/a |
| | 2 | f | 60~80 | 3 | Liver zirrhosis with esophageal varices | 1050 | n/a | skull+, meninges-, brain n/a |
| | 3 | f | 40~60 | 8 | Ethanol dependency, Gastritis, Hypothyreosis | 1140 | 6 | skull-, meninges-, brain n/a |
| | 4 | m | 40~60 | 7 | Arterial Hypertension, Arterial disease, Adipositas | 1570 | 14 | skull+, meninges-, brain n/a |
| | 5 | m | 80~100 | 3 | Arteriosclerosis, coronary artery disease (3VD), COPD | 950 | 12 | skull-, meninges n/a, brain n/a |
| | 6 | m | 40~60 | 3 | CI, IHD, DM | 1535 | 9 | skull-, meninges n/a, brain n/a |
| | 7 | f | 80~100 | 5 | Emphysema, IHD, CI, CKD | 1030 | n/a | n/a |
| | 8 | m | 60~80 | 6 | Metastased rectal CA, mitral valve sclerosis | 1390 | n/a | skull-, meninges n/a, brain n/a |
| | 9 | m | 40~60 | 6 | liver cirrhosis, esophagus vericosis, CKD, emphysema | 1500 | 1 | skull-, meninges n/a, brain n/a |
| | 10 | m | 80~100 | 4 | asbestosis, IHD, bronchial CA, | 1385 | n/a | skull-, meninges n/a, brain n/a |
| | 11 | f | 80~100 | 3 | Colon-CA, Apoplex, dementia, IHD | 1290 | n/a | skull-, meninges n/a, brain n/a |
| | 12 | m | 60~80 | 3 | n/a | 1350 | n/a | skull+, meninges-, brain n/a |
| | 13 | f | 60~80 | 4 | DM | 1085 | n/a | skull+, meninges+, brain- |
| | 14 | f | 60~80 | 2 | JAK2 pos. Polycythaemia vera, art. Hypertonie | 1380 | 55 | skull+, meninges+, brain- |
| | 15 | m | 60~80 | 8 | chronic kidney insufficiency,<br>prostate carcinoma, arterial hypertension | 1632 | 34 | skull+, meninges+, brain- |
| | 16 | f | 80~100 | 3 | breast cancer | 1210 | n/a | skull+, meninges-, brain- |
| | 17 | f | 60~80 | 12 | Alzheimer's type dementia, cardiac arrhythmia,<br>high blood pressure, chronic cardiac insufficiency | 1365 | 3 | skull+, meninges+, brain- |
| COVID-19 brain cortex | | | | | | | | |
| | 1 | f | 60~80 | n/a | n/a | n/a | n/a | skull n/a, meninges n/a, brain- |
| | 2 | m | 40~60 | 4 | n/a | 1698 | n/a | skull n/a, meninges n/a, brain- |
| | 3 | n/a | n/a | n/a | n/a | n/a | n/a | skull n/a, meninges n/a, brain- |
| | 4 | n/a | n/a | n/a | n/a | n/a | n/a | skull n/a, meninges n/a, brain- |
| | 5 | n/a | n/a | n/a | n/a | n/a | n/a | skull n/a, meninges n/a, brain- |
| | 6 | n/a | n/a | n/a | n/a | n/a | n/a | skull n/a, meninges n/a, brain- |
| | 7 | n/a | n/a | n/a | n/a | n/a | n/a | skull n/a, meninges n/a, brain- |
| | 8 | m | 60~80 | 1 | n/a | n/a | n/a | skull n/a, meninges n/a, brain- |
| | 9 | m | 80~100 | n/a | n/a | n/a | n/a | skull n/a, meninges n/a, brain- |
| | 10 | n/a | n/a | 2 | Essential hypertension; chronic lower respiratory diseases;<br>Alzheimer's disease | 1045 | n/a | skull n/a, meninges+, brain- |
| | 11 | n/a | n/a | n/a | n/a | n/a | n/a | skull n/a, meninges+, brain- |
| Control skull/meninges | | | | | | | | |
| | 1 | m | 80~100 | 1 | intestinal infections; Atherosclerosis; Heart failure | 1268 | none | |
| | 2 | m | 60~80 | 2 | chronic ischemic heart disease; Essential hypertension | 1270 | none | |
| | 3 | m | 80~100 | 2 | nutrition/metabolic; mental/behavioral disorders;<br>Renal failure; heart disease | 1183 | none | |
| | 4 | m | 80~100 | 2 | Malignant neoplasm of prostate | 1204 | none | |
| | 5 | n/a | n/a | n/a | n/a | n/a | none | |
| | 6 | n/a | n/a | n/a | n/a | n/a | none | |
| | 7 | n/a | n/a | n/a | n/a | n/a | none | |
| | 8 | n/a | n/a | n/a | n/a | n/a | none | |
| | 9 | n/a | n/a | n/a | n/a | n/a | none | |
| | 10 | n/a | n/a | n/a | n/a | n/a | none | |
| Control brain cortex | | | | | | | | |
| | 1 | m | 40~60 | 2 | n/a | n/a | none | |
| | 2 | m | 40~60 | 5 | n/a | n/a | none | |
| | 3 | m | 0~20 | 3 | n/a | n/a | none | |
| | 4 | m | 60~80 | 8 | n/a | n/a | none | |
| | 5 | f | 80~100 | 5 | n/a | n/a | none | |
| | 6 | f | 20~40 | 1 | n/a | n/a | none | |
| | 7 | f | 80~100 | 3 | Mental/behavioral disorders; Essential hypertension | 1190 | none | |
| | 8 | f | 80~100 | n/a | n/a | 1150 | none | |
| Patients died between 2021-2022 | | | | | | | | |
| | 1 | m | 40~60 | n/a | | | | |
| | 2 | m | 60~80 | 1 | | | | |
| | 3 | m | 80~100 | 1 | | | | |
| | 4 | f | 20~40 | 1 | | | | |
| | 5 | m | 60~80 | n/a | | | | |
| | 6 | m | 80~100 | 10 | | | | |
| | 7 | m | 60~80 | 6 | | | | |
| | 8 | m | 40~60 | n/a | | | | |
| | 9 | f | 60~80 | n/a | | | | |
| | 10 | m | 60~80 | n/a | | | | |
| | 11 | m | 40~60 | 1 | | | | |
| | 12 | m | 40~60 | 1 | | | | |
| | 13 | m | 80~100 | 1 | | | | |
| | 14 | f | 60~80 | 1 | | | | |
| | 15 | f | 60~80 | 1 | | | | |
| | 16 | m | 60~80 | 1 | | | | |
| | 17 | m | 80~100 | n/a | | | | |
| | 18 | m | 40~60 | n/a | | | | |
| | 19 | f | 40~60 | n/a | | | | |
| | 20 | f | 60~80 | 13 | | | | |
| | 21 | m | 40~60 | 11 | | | | |
| | 22 | f | 80~100 | 12 | | | | |
| | 23 | m | 60~80 | 10 | | | | |
| | 24 | m | 40~60 | n/a | | | | |
| | 25 | m | 80~100 | n/a | | | | |
| | 26 | f | 80~100 | n/a | | | | |
| | 27 | m | 60~80 | n/a | | | | |
| | 28 | f | 80~100 | n/a | | | | |
| | 29 | f | 80~100 | n/a | | | | |
| | 30 | f | 20~40 | n/a | | | | |
| | 31 | n/a | n/a | n/a | | | | |
| | 32 | n/a | n/a | n/a | | | | |
| | 33 | m | 60~80 | 19 | | | | |
| | 34 | m | 20~40 | 1 | | | | |
| n/a: not available | | | | | | | | |
| PMI: postmortem interval (days) | | | | | | | | |
| onary disease, DM: diabetes mellitus, IHD: ischemic heart disease, RI: renal insufficiency, Tx: Transplantation | | | | | | | | |
